# Effects of microplastic ingestion on hydrogen production and microbiomes in the gut of the terrestrial isopod *Porcellio scaber*

**DOI:** 10.1101/2022.06.22.497054

**Authors:** Linda Hink, Anja Holzinger, Tobias Sandfeld, Alfons R. Weig, Andreas Schramm, Heike Feldhaar, Marcus A. Horn

## Abstract

Microplastic (MP) pollution is an environmental burden. MP enters food webs via ingestion by macrofauna, including isopods (*Porcellio scaber*) in terrestrial ecosystems. However, MP-effects on the host and its gut microbiome are largely unknown. We tested the hypothesis that biodegradable (polylactic acid, PLA) and non-biodegradable (polyethylene terephthalate, PET; polystyrene, PS) MP have contrasting effects on *P. scaber* mediated by changes of the associated gut microbiome. Although the isopods avoided food containing PS, isopod fitness after eight-week MP-exposure was unaffected. Qualitative and quantitative 16S rRNA gene and 16S rRNA analyses of gut microbiomes indicated general MP effects, MP-type specific indicator taxa, and stimulation by PLA compared to MP-free controls. Isopods emitted hydrogen, and its production increased and decreased after PLA-food and PET- or PS-food ingestion, respectively, relative to controls as indicated by microsensor measurements. Gut pH was unaffected by MP. We identified the gut of *P. scaber* as significant mobile source of reductant for soil microbiomes likely due to *Enterobacteriaceae* related fermentation activities that were stimulated by lactate generated during PLA-degradation. The findings suggest negative effects of PET and PS on gut fermentation, modulation of isopod hydrogen emissions by MP pollution, and the potential of MP to affect terrestrial food webs.

## Introduction

Today’s modern life without plastic is inconceivable, as plastic is highly versatile and can be applied in various sectors ranging from the packaging to the building and construction sector with a likewise wide range of lifetimes from less than a year to several decades. To date more than 8.3 billion tonnes have been produced, out of which about 60% have been discarded. A large proportion of the plastic waste is either disposed of in landfills or ends as litter in the natural environment [1]. Hence plastic is not only advantageous, but also has become a ubiquitous man-made environmental burden. Due to weathering and fragmentation, large plastic items are gradually transformed to slowly degrading microplastic (MP) smaller than 5 mm [2, 3]. Although biodegradation by specialised microorganisms originating from plastic polluted sites, landfills or animal intestines has been observed [4], MP only degrade slowly in natural environments and persist over decades [5]. While most attention so far focused on oceanic environments, the annual plastic release to terrestrial environments is 4-23 times higher mostly due to agricultural practices and littering [6]. Conventional plastics, such as polyethylene (PE), polyethylene terephthalate (PET), polystyrene (PS) and polyvinyl chloride (PVC), are considered to be extremely resistant to biodegradation. Biodegradable polymers such as polylactic acid (PLA) and polycaprolactone (PCL) are therefore becoming popular as an alternative. However, their degradation is incomplete and slow under environmental conditions [7].

MP may not only alter the physical and chemical soil properties, but may also affect the soil biota [8]. Unintentional ingestion of MP by soil-dwelling and especially soil-feeding macrofauna is likely and can have negative fitness effects on organisms like earthworms, nematodes or collembola [9, 10]. Subsequently MP may be transferred to higher trophic levels as shown for chickens that acquired MP from soil by feeding on earthworms [11].

Alike earthworms, also terrestrial isopods (i.e., woodlice - Isopoda - Oniscidea) are widespread decomposers with a density that can exceed 1000 individuals m^-2^ [12]. These organisms are sensitive to contaminants, e.g. pesticides or heavy metals, and thus are suited for soil ecotoxicity testing in laboratory and field bioindicator studies [12, 13]. As soil-dwellers they mainly feed on decaying leaf litter and wood, and it has been shown that weathered feed colonized by microbes is actually favoured [14, 15]. Their presence in soil enhances soil nutrient cycling due to fragmentation and transportation of organic material along with microorganisms [16]. Hence, it can be considered that they likewise contribute to the fragmentation and transportation of accidently ingested MP and even distribution of pathogenic microorganisms as plastic surfaces are suitable for colonization [17–19]. Many studies have investigated the effects of pollutants on the isopod *Porcellio scaber* (common rough woodlouse) [13]. However, to date there are only a few studies investigating life history traits after MP ingestion: No or only minor effects on survival, feeding rate, body mass or energy reserves in the digestive glands were obtained for isopods exposed to PE particles, tire particles or polyester fibers [20–22]. However, this does not necessarily mean that *P. scaber* is only marginally affected by MP ingestion. The effects are possibly less obvious after relatively short exposure of only a few weeks and the detection of sublethal effects requires assessment of other parameters. Accordingly, immune response parameters are affected by MP (polyester fibers and tire particles) ingestion [23, 24]. Such effects are potentially linked to effects on the gut microbiome, but the effects of MP exposure on the gut microbiome of *P. scaber* have not been investigated yet.

Generally, an intact gut microbiome is important for the development, nutrition and immunity and this also applies for isopods that possess a more dynamic microbiome compared to higher organisms, such as mammals [25]. The nutrient content in the common diet of isopods (leaf litter and dead wood) is generally very low and it is suggested that isopods rely on microbes colonizing and degrading the decaying plant material and thereby providing nutrients for the host, or the microbes themselves are digested and serve as nutrient-rich source [15, 26, 27]. With a reduced microbial cell number, it is likely that the latter applies for the anterior section of the hindgut, while microbes proliferate towards the posterior section, where mainly anaerobic conditions prevail [28–30]. The most abundant groups in *P. scaber’s* gut microbiome have been assigned to Proteobacteria, Bacteroidota, and Actinobacteria commonly inhabiting insect intestines [26, 31–33]. All of these phyla contain members that possess a facultative or even obligate anaerobic lifestyle and therefore, fermentative microbes including hydrogen producers may play an important role as shown for other invertebrates (e.g., termites and earthworms) [34, 35]. However, whether or not fermentation is an ongoing process in the gut of *P. scaber* remains to be determined.

The gut microbial community and hence the digestive processes can indeed be modulated by MP ingestion as shown for several soil invertebrates: In the gut of mealworms (*Tenebrio molitor* larvae), PE and PS can be degraded with strong association of species within the *Enterobacteriaceae* [36]. Adverse effects of MP ingestion on life history traits (e.g., reduced growth and reproduction rates) appear along with alteration of the gut microbiome of springtails (*Folsomia candida;* ingestion of PVC or PE) [37, 38] and potworms (*Enchytraeus crypticus;* ingestion of PS) [39]. Studies of *P. scaber’s* gut microbiome after MP ingestion are lacking to date.

This study aims to investigate the effects of conventional non-biodegradable MP particles, PET and PS, and biodegradable PLA on *P. scaber* with respect to fitness, gut microbiome and fermentation potential in the gut, with the underlying hypothesis that biodegradable and non-biodegradable MP have contrasting effects on *P. scaber* mediated by changes of the associated gut microbiome. This was tested in MP-feeding experiments, microsensor analysis of prevailing conditions in the gut with respect to pH, oxygen and hydrogen (as a measure for ongoing microbial fermentation) concentration and analysis of the gut microbiome. In addition, the food microbiome was analysed to investigate its influence on the gut microbiome.

## Materials and Methods

### Food preparation and isopod collection

Food pellets consisting of withered leaves (mainly maple leaves; 42%), ground commercial rabbit food (25%) and potato powder (33%) were prepared as described in Žižek *et al*. [40]. For the pellets that additionally contained MP particles, 2.5% or 5% (w/w) PLA (NatureWorks, Naarden, The Netherlands), PET (Veolia, Berlin, Germany) or PS (Ineos Styrolution, Ludwigshafen, Germany) was added to the mixture. Granules were ground to fragments using a cryo ball mill (Retsch, CryoMill, Germany) followed by sieving to obtain fragments ranging from 75-150 μm in diameter of irregular shape prior usage.

*P. scaber* individuals (only adults; weight >30 mg) as model isopods were collected in a garden near the campus of the University of Bayreuth (Germany) or the Leibniz University of Hannover (Germany) between February and May 2020. The animals were kept in boxes (40 cm x 30 cm x 25 cm) filled with damp soil, leaves, and tree bark prior performance of independent experiments assessing hydrogen and methane emission rates of whole isopods, microsensor profiles of pH, hydrogen and oxygen concentrations of isopod guts, bacterial community composition of the isopod guts and food pellets, as well as fitness effects and food choice (for the latter see Supplementary Material and Methods).

### Molecular hydrogen, oxygen and pH microsensor measurements from isopod guts

Microsensor measurements were performed to identify the location and level of hydrogen production within isopods as an estimate for the fermentation potential and to assess MP effects on the conditions inside the gut of the isopods. 1 g of food pellets containing no MP or 5% PLA, PET or PS were mixed with 2 ml 1% agar (∼60°C), spread on a petri dish and cooled to room temperature. Twelve isopods per treatment were placed on these petri dishes and kept at room temperature in the dark. The food was exchanged after 3 days and isopods were kept for 3 further days. Prior to gut dissection, the isopods were placed on ice for several minutes in order to lower their mobility. Each gut was embedded within a small glass chamber in 1% low-melt agarose in insect Ringer’s solution. Coverslips and microscope slides (7.5 x 2.5 x 0.1 cm) were used for construction of chambers similar to those in Brune *et al*. [41]: A coverslip at the bottom of the chamber and two microscope slides on top of each other were arranged to each side of the chamber providing the dimensions of 2.5 cm length, 1.0 cm width and 0.2 cm depth. The bottom of the chamber was filled with a layer of molten 1% low melt agarose in insect Ringer’s solution and after solidification a freshly dissected full isopod gut was placed on top of it. Then a top agarose layer (not warmer than 40°C) was cast in the chamber, which was immediately covered with a coverslip before solidification. The embedded gut was placed on another 2-mm thick agarose bed in a weighing boat and covered with insect Ringer’s solution.

Custom made microsensors for oxygen, hydrogen and pH [34, 42–44] with tip diameters <20 µm were used for recording radial profiles of the anterior (at a distance of 1 mm from the front end), the median and the posterior (at a distance of 1 mm from the rear end) of isopod guts (see positions in Fig. S1). Measurements were performed at room temperature. For pH measurements, a bridge consisting of a syringe barrel filled with the same agarose as the agarose bed was constructed between this bed and a Red Rod reference electrode. The sensors were connected to a four-channel multimeter with a built-in 16-bit A/D converter (Unisense Microsensor Multimeter, Ver 2.01; Unisense A/S, Denmark). Pre-polarized sensors were calibrated prior gut profile measurements: a two-point calibration with a 0.7 M alkaline ascorbate solution (0 µM oxygen) and an air-saturated Ringer’s solution (265.6 µM oxygen) for the oxygen microsensor; a three-point calibration with pH 4, 7 and 10 buffers for the pH microsensor; a calibration with multiple points ranging between 0 and 50 µM for the hydrogen microsensor. Data acquisition and control of the micro-profiling system was enabled with the software program SensorTrace PRO (Unisense A/S, Denmark). Profiling through the guts was performed with 50 µm spatial resolution. Oxygen and pH sensors were allowed to equilibrate for 5 s prior data acquisition. For hydrogen sensors, 10 s equilibration time were required. The measurements with the different sensors were performed with different guts. Three guts per treatment and microsensor were analysed at three positions each (anterior, median and posterior).

### Determination of molecular hydrogen and methane emission from whole isopods

In order to confirm hydrogen emissions under *in vivo* conditions, hydrogen production rates of whole isopods were determined. In addition to hydrogen, methane is relevant in the anaerobic food chain and was also analyzed. Therefore, adult isopods were collected and surface sterilized with 70% ethanol. These animals were not subjected to a MP-treatment. Groups of three individuals each were placed in three 3-mL Exetainer (Labco, Lampeter, UK). The vials were sealed with airtight lids (caps with butyl septa) and overpressure was applied to all vials via injection of 2 ml air. Headspace hydrogen and methane mixing ratios were analysed for a period of 10 h with a gas chromatograph coupled to a pulsed discharge helium ionization detector as described (7890B, Agilent Technologies, JAS GC systems, Moers, Germany) [45].

### Isopod feeding experiment with assessment of fitness parameters

Groups of 10 isopods (7 females, 3 males) were kept in glass jars (diameter: 10.8 cm; volume: 370 ml) with the bottom covered with moist filter paper in a climate chamber with a 16 h light and 8 h dark circle at 16°C and 85% humidity. Isopod groups were exposed to food pellets without or with 2.5% or 5% PLA, PET or PS for eight weeks. Each treatment was performed in 5 replicates. One food pellet was placed in each jar and exchanged every other day. At the same time survival of isopods was checked and dead individuals were removed. Once a week the glass jars were cleaned and the filter papers were replaced. Locomotor activity tests were performed after two, four and six weeks (see Supplementary Material and Methods). After eight weeks, the isopods were weighed and the percentage weight gain to the initial weight was calculated. Further, one gut per replicate was dissected and these guts as well as the food pellet of the respective treatment were frozen in liquid nitrogen and kept at -80°C prior microbiome analysis.

### Nucleic acid extraction, DNase treatment and reverse transcription

Prior extraction of DNA and RNA the weight of the whole isopod guts was determined. The nucleic acids were also extracted from subsamples of the food pellets (∼50 mg). The extraction protocol was performed according to Griffiths *et al.* [46] with some modifications: i) 2-ml screw cap tubes used during initial cell lysis were filled with Ø0.1-mm and Ø0.5-mm zirconium beads, 150 mg each, and one Ø3-mm glass bead; ii) cell lysis by bead beating for 30 s at 5.0 m s^-1^ was performed twice with an intermitted cooling on ice for 30 s; iii) nucleic acids were precipitated on ice for 2 h; iv) air-dried nucleic acid pellets derived from gut and food samples were resuspended in 30 μl and 60 µl RNase-free water, respectively. Verification of nucleic acid extracts was assessed via agarose gel electrophoresis and spectrophotometric measurements. DNase treatment was applied to a 13-µl subsample of each extract using the TURBO DNA-free™ Kit (Invitrogen, Thermo Fisher Scientific, Waltham, MA, USA) according to the manufacturer’s instructions. Reverse transcription of 10 µl RNA was performed using LunaScript RT Supermix Kit (New England Biolabs, Ipswich, MA, USA) according to the manufacturer’s protocol. Negative controls without template RNA (water treated with DNase) and, for each sample, without reverse transcriptase were performed. Samples were kept on ice for further processing.

### Analysis of bacterial 16S rRNA genes and 16S rRNA

Bacterial 16S rRNA genes and 16S rRNA (after reverse transcription) were quantified by qPCR using primers Bact_341F and Bact_805R [47]. Each sample was analyzed in duplicate 10-µl reactions containing 5 µl Luna Universal qPCR Master Mix (New England Biolabs, Ipswich, MA, USA), 2 µM of each primer, 10 µg bovine serum albumin and 2 µl of 1:100 diluted template (c)DNA. Negative controls contained sterile water instead of template (c)DNA. Standards consisted of serially diluted (10^2^ to 10^6^ gene copies per µl) M13uni/rev PCR products of a pGEM-T vector with a 16S rRNA gene inserted. The amplification was performed in a CFX Connect Real-Time PCR System (Bio-Rad, Feldkirchen, Germany) with the following cycling conditions: 2 min at 95°C, 35 cycles of 30 s at 95°C, 50 s at 60°C (combined annealing and elongation) and a plate read after 10 s at 80°C. The amplification efficiencies ranged between 96% and 98%, the r^2^ values were ≥0.99. The specificity of the amplification was verified by melting curve analysis (from 60°C to 95°C in 0.5°C-intervals for 5 s each) and agarose gel electrophoresis. Specific amplification of archaeal 16S rRNA (using primers A519F [48] and A1017R [49]) and [Fe-Fe]-hydrogenase genes (encoding for enzymes that catalyze the production of hydrogen in obligate anaerobic fermenting bacteria, using primers from Schmidt *et al.* [50] and Xing *et al*. [51]) was tested for several gut and food samples, but consistently failed.

Bacterial 16S rRNA genes and 16S rRNA were also analyzed by high-throughput amplicon sequencing. Amplicons were generated using the same primer pairs as for the qPCR, but tagged with specific adapters (Illumina, San Diego, USA). Each reaction mixture contained 12.5 µl Kapa HiFi HotStart ReadyMix (Roche, Mannheim, Germany), 0.5 µM of each primer, 5 µg bovine serum albumin and 2.5 µl of 1:10 diluted template (c)DNA. Sterile water, instead of template was applied for negative controls. The PCR was performed in a Thermocycler (Biozym Scientific GmbH, Hessisch Oldendorf, Deutschland) with 3 min initial denaturation at 95°C, followed by 30 cycles with 20 s at 98°C, 15 s at 55°C and 15 s at 72°C, and an end-elongation step at 72°C for 1 min. Specificity of the amplification was confirmed by agarose gel electrophoresis. The amplicons were purified using the GeneRead size selection kit (Qiagen, Hilden, Germany). Library preparation was performed using the Nextera XT Index Kit (Illumina, San Diego, United States) as given elsewhere [52] and the Illumina MiSeq version 3 chemistry was applied for 2×300 bp paired-end sequencing. Details regarding generation of amplicon sequencing variants (ASVs) and further analyses of the sequences are provided in the Supplementary Material and Methods.

### Statistical analysis

All statistical analyses were conducted using the statistical platform R version 4.1.1 [53]. Differences in weight gain, food choice and speed index of the isopods as well as maximum hydrogen concentration and minimum pH in the isopod guts were analyzed by fitting linear models, performing ANOVA tests and Tukey *post hoc* comparisons using the *multcomp* package [54]. The survival probability of isopods and confidence intervals were calculated using the *survival* package [55]. Abundance data obtained from qPCR analysis was investigated by a factorial two-way ANOVA (MP-treatment and percentage as factors). If the initial model has not met normality assumptions, the data was transformed using the transformTukey function in the *rcompanion* package [56], and then the adjusted model met the required assumptions. Tukey *post hoc* tests were applied to assess significant differences in means. Whether the community compositions differed between the treatments was assessed with PERMANOVA tests and pairwise comparisons applied on the Aitchison distance matrix using the adonis2 and the adonis.pair function in the *vegan* [57] and *EcolUtils* [58] packages of R, respectively.

## Results

### Effects of microplastic on isopods

When *P. scaber* had the choice between food containing no MP or 5% PLA, PET or PS, a significant avoidance of PS-food was observed (Fig. S2a). However, when there was no choice given, any food was ingested. Neither the survival (∼20% dead individuals after eight weeks; Fig. S3), nor the weight gain was affected by the MP-diet (p = 0.556; Fig. S2b). The speed index for isopods exposed to 2.5% PET was significantly lower than that of those exposed to 2.5% PS, nevertheless none of the MP-food had a significant effect compared to the control-food (Fig. S2c).

### Physicochemical conditions in the gut of isopods and effects of microplastic ingestion

Radial oxygen micro-profiles revealed anoxic conditions at any position in the guts of isopods fed with any diet, and pH ranged from 5 to 7 (Figs. S4, S5). Minimum gut pH values were not affected by the MP-treatment nor an interaction of position and MP-treatment (ANOVA; p > 0.05). The minimum pH was significantly more acidic in the anterior (pH 5.2) than in the median to posterior (pH 5.8) positions (Fig. S5, Table S1). Hydrogen was highest inside the isopod guts at all measured positions (Figs. 1, S6). Hydrogen concentrations of up to ∼20 µM were detected in the gut center of isopods fed with MP-free control food. Isopods fed with PLA-food showed highest and those fed with PET- or PS-food lowest gut hydrogen concentrations. At the gut median, maximum hydrogen concentrations of ∼30 µM were significantly higher in isopods fed with PLA-food than with other food (Table. S2). At the posterior position of the gut, maximum hydrogen concentrations of ∼5 µM were significantly lower in isopods fed with PET- and PS-food than with control- and PLA-food. It is also worth mentioning that in guts of isopods fed with control- and PLA-food, the hydrogen formation activity was significantly higher towards the posterior than in the anterior end.

**Fig. 1:**
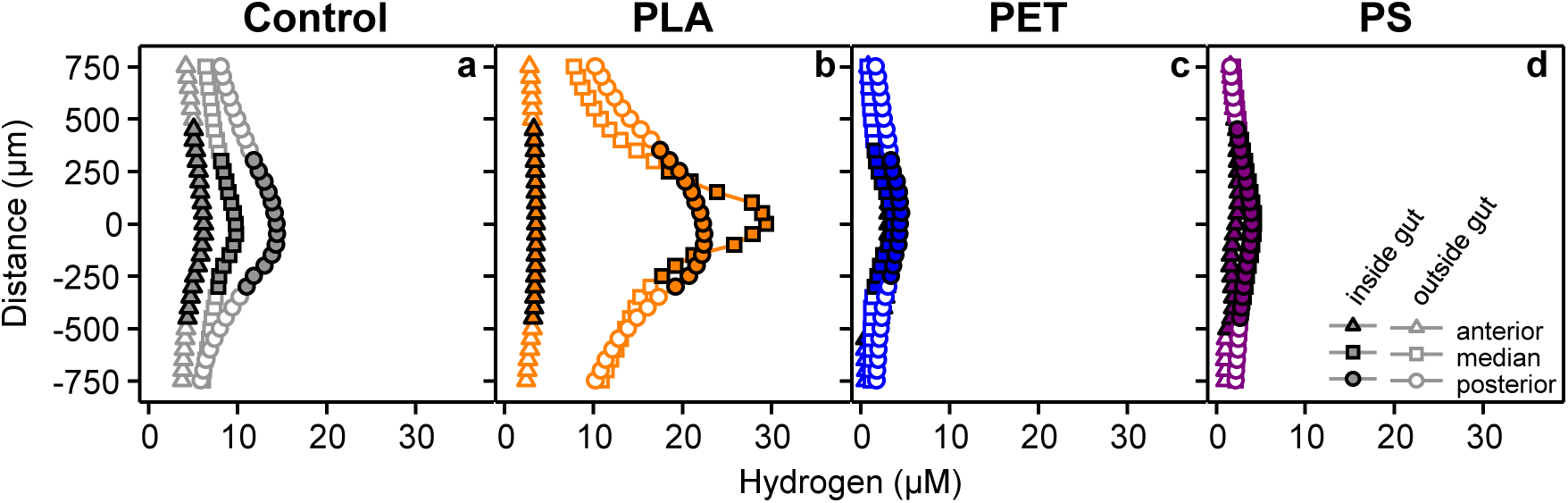
Representative radial hydrogen profiles of isopod guts. The isopods were fed with food containing no microplastic particles (control; **a**) or 5% PLA (**b**), PET (**c**) or PS (**d**) for 6 days prior gut extraction and subsequent embedding in agarose and microsensor measurements. For each gut, profiles were recorded from the anterior, median and posterior. Analyses of two more guts per treatment are displayed in Fig. S6. Closed and open symbols represent measured concentrations inside and outside (in agarose) the guts, respectively. The distance of 0 µm indicates the center of the tube-like gut.

### Hydrogen emission potential of whole isopods

Hydrogen mixing ratios in the headspace of vials containing whole isopods (analyzed directly after collection and not subjected to MP-treatments) increased linearly over time without appreciable delay (Fig. 2), demonstrating that hydrogen is indeed emitted from whole isopods under *in vivo* conditions. The emission rates were highly variable with an average rate of 0.83 ± 0.51 ng hydrogen isopod^-1^ h^-1^ resulting in a final amount between 1 and 24 ng hydrogen isopod^-1^ after 10 hours of incubation.

**Fig. 2:**
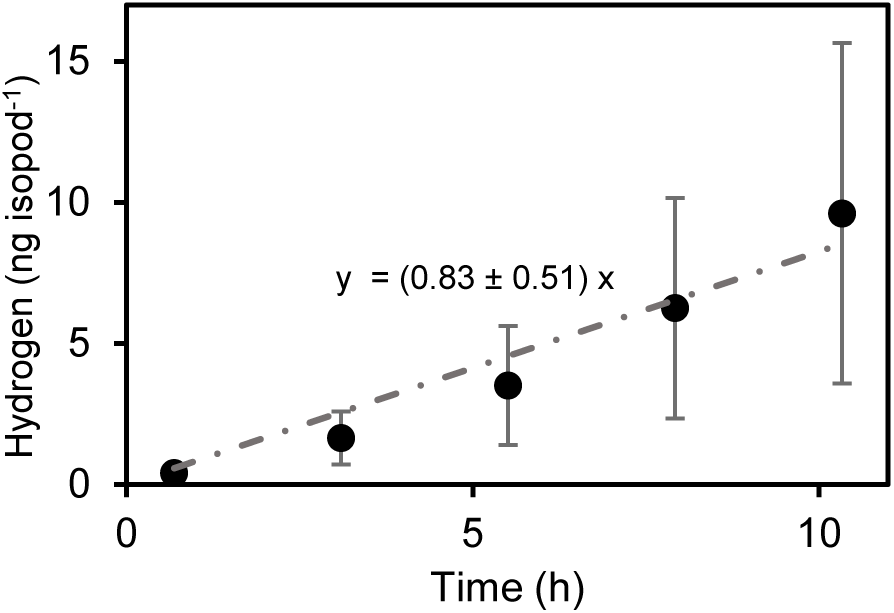
Hydrogen accumulation in the headspace of whole isopods incubated under air for 10 h. Values represent means of triplicate vials each containing three isopods that were analyzed directly after collection and not subjected to a MP-exposure experiment. The dot-dashed line indicates a linear regression of the hydrogen mixing ratios. Means and standard errors of three replicates are plotted. The regression equation including the standard error of the slope is shown near the regression line.

### Impact of microplastic on 16S rRNA gene and 16S rRNA abundance in food and guts of isopods

Bacterial 16S rRNA abundances were essentially one order of magnitude higher in the isopod guts than in the food pellets, while the opposite applied for 16S rRNA genes (Fig. 3). Such findings were supported by the 16S rRNA:16S rRNA gene ratios that were consistently higher in the guts than in the food despite a high variability among replicates (Fig. 3c,f). 16S rRNA gene abundances obtained from the guts were significantly higher in isopods exposed to PLA-food compared to the other treatments (Fig. 3a). A similar stimulation was reflected at 16S rRNA level, but not in 16S rRNA:16S rRNA gene ratios (Fig. 3b,c). MP had no significant effects on the 16S rRNA gene and 16S rRNA abundances in the food pellets and ratios thereof (Fig. 3d,e,f).

**Fig. 3:**
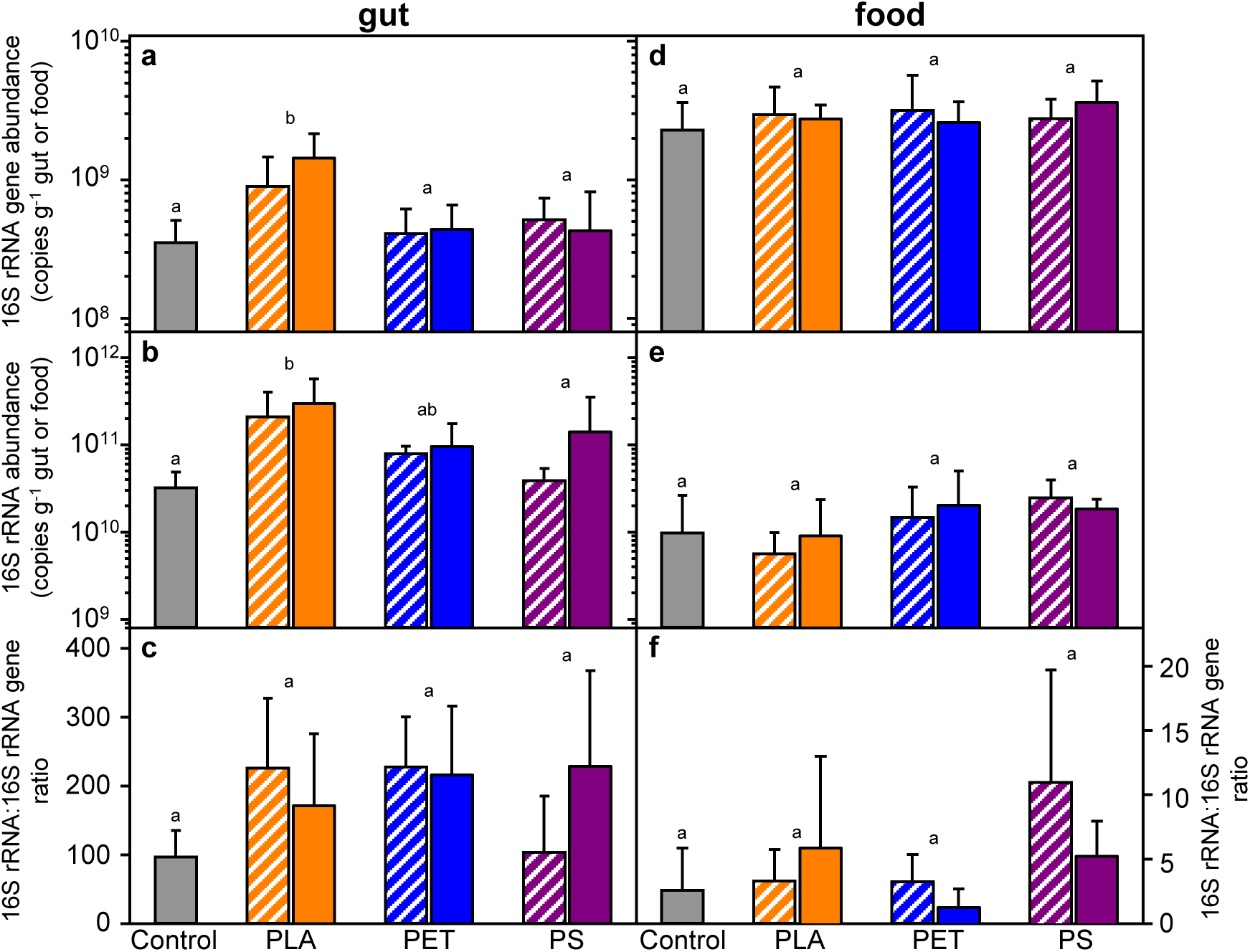
Effects of MP ingestion on the abundance of bacterial 16S rRNA genes and 16S rRNA in the gut and respective food of *P. scaber.* Nucleic acid extracts derived from the guts of isopods (**a, b, c**) that were exposed to control- or 2.5%-MP- (striped) or 5%-MP- (no pattern) food pellets (**d, e, f**) were directly used for quantification of genes (**a, d**). A subsample of each extract was subjected to DNase treatment and subsequent reverse transcription for analysis of 16S rRNA (**b, e**). In addition, the 16SrRNA:16S rRNA gene ratios were calculated (**c, f**). The data was corrected for the proportion on endosymbionts obtained from sequencing analysis (see supplementary Fig. S7 for comparison of uncorrected and corrected data). Means and standard deviations of five replicates are plotted. Statistical analysis revealed no effect of the concentration of MP applied and therefore, significant differences in means indicated by different lower letters above the bars are related the MP treatment regardless the dosage.

### Impact of microplastic on the bacterial communities

The bacterial communities in the isopod guts were highly diverse (Fig. S8c,d). Highest relative gene abundances were found for taxa within the Actinobacteria (mainly *Microbacteriaceae*), Bacteroida (mainly *Flavobacteriaceae*), Gammaproteobacteria (mainly *Enterobacteriaceae* and *Vibrionaceae*) and Verrucomicrobiae (mainly *Opitutaceae*) (Fig. S8c). All of these taxa, but the Actinobacteria to a minor extent, were also found at 16S rRNA level (Fig. S8d).

Gut communities differed from the food communities as indicated by principal coordinates analysis (PCoA; PERMANOVA; p < 0.005; Fig. S9a; for further details see Supplementary Results). ASVs assigned to *Vibrio rumoiensis* correlated well with gut communities. To elucidate the effects of MP, the datasets were analyzed separately for the gut and the food communities (Figs. 4, S9b). In addition to *Vibrio rumoiensis*, the gut communities were mainly affected by *Enterobacteriaceae* and *Microbacteriaceae* (Fig. 4). PERMANOVA analysis revealed that gut communities differed significantly due to MP treatment. An effect of MP treatment and dosage was obtained. For the latter, significant difference between communities exposed to 2.5% and 5% MP has been confirmed by a pairwise comparison (Table S3). Such pairwise comparisons essentially confirmed differences between gut communities of isopods fed with PLA-food and those fed with PET- or PS-food with low p-values (p < 0.07), but failed to confirm other MP treatment effects (p > 0.15) (Table S4).

**Fig. 4:**
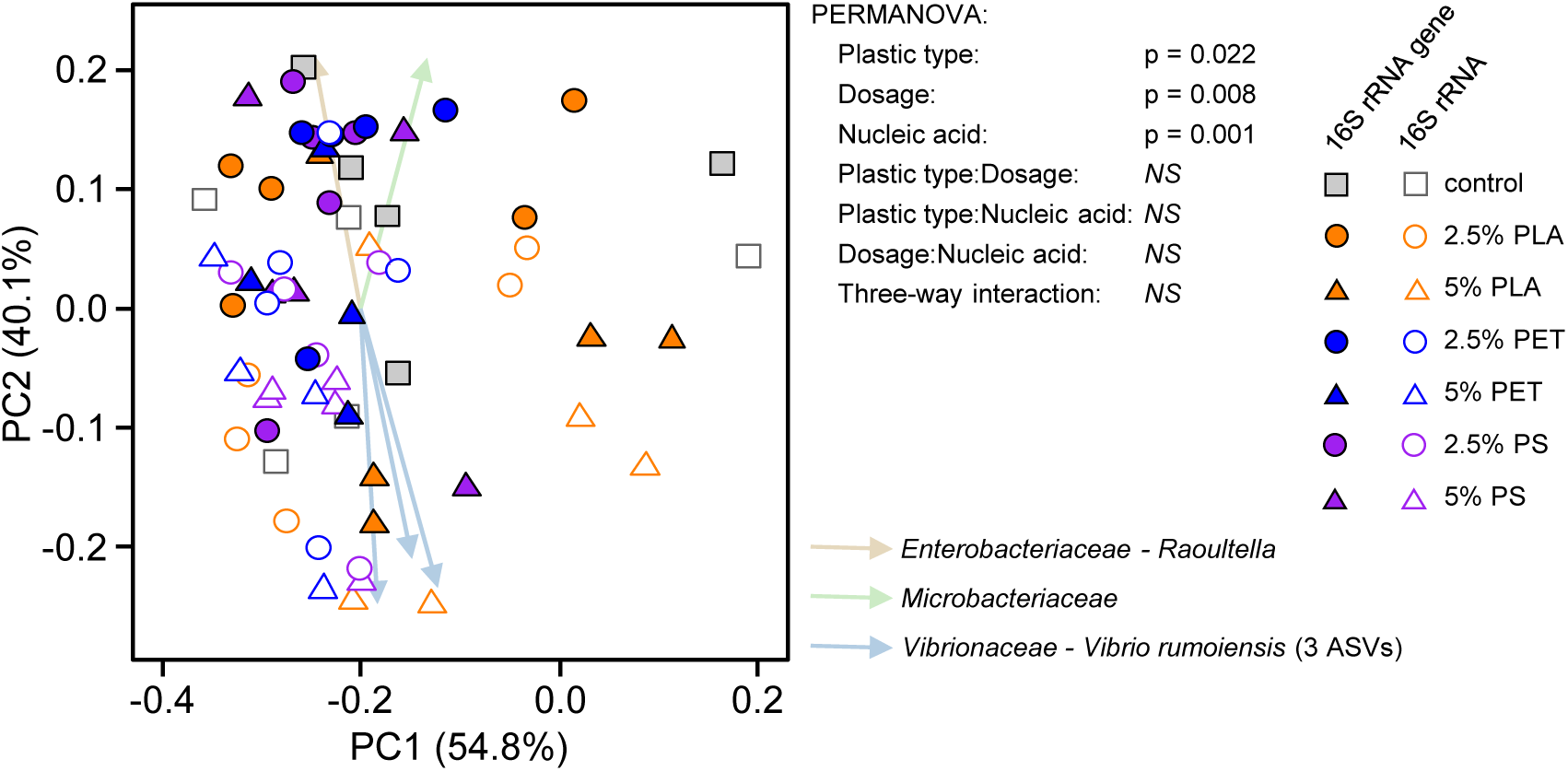
Beta diversity of the active bacterial gut communities. PCoA plots are based on Aitchison distance matrixes derived from analyses of the 16S rRNA genes and 16S rRNA. Results of the PERMANOVA analyses are given for each plot. Arrows represent ASVs assigned on family and genus/species level (if applicable) that were highly correlated with the separation of samples.

### Shared, unique and indicator taxa with respect to MP-treatments

Most genera were shared among all treatments in isopod guts (43-49%; Figs. 5, S10a). On 16S rRNA gene and 16S rRNA level, only a few genera were uniquely found in guts of isopods fed with PET- or PS-food (2-4%), but more in case of guts of isopods fed with control- or PLA-food (7-13%). Moreover, more genera in guts of isopods exposed to control-food were shared with those exposed to PLA-food (8-13%) than with those exposed to PET- (1%) or PS-food (0%). Genera within *Alcaligenaceae* were among exclusive taxa in guts of PLA-food exposed isopods.

**Fig. 5:**
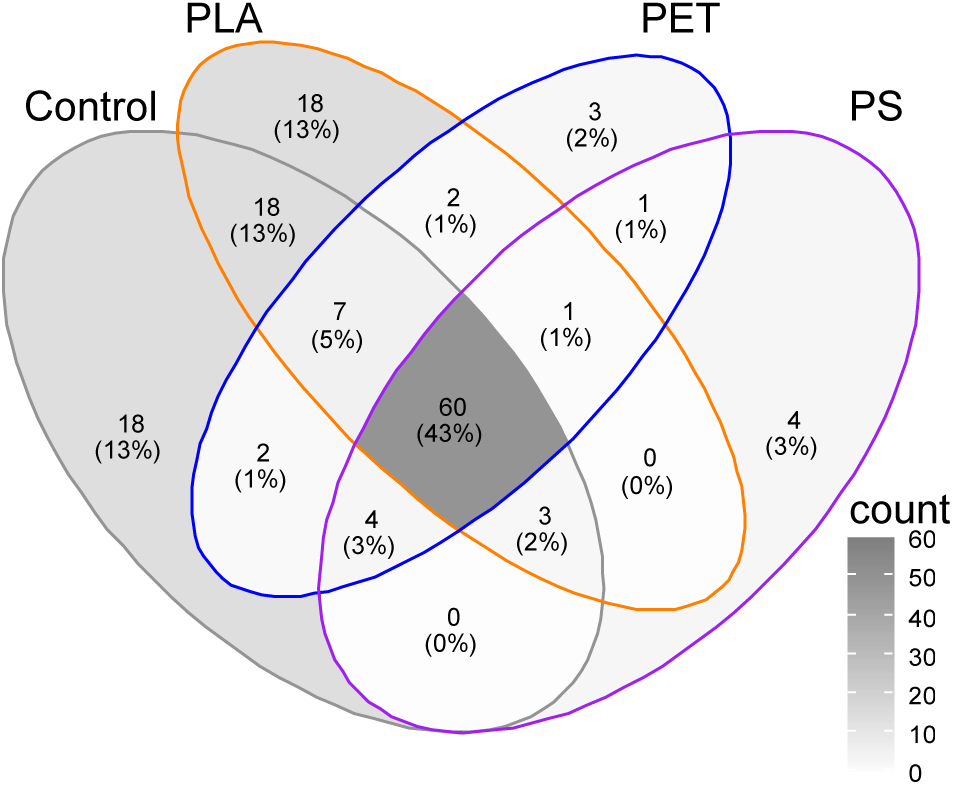
Shared and unique numbers and proportions of taxa among guts of isopods fed with control-, PLA-, PET- and PS-food on 16S rRNA level. Only taxa on genus level (if applicable) that occur in at least 30% of the replicates were included for the calculation of the Venn diagram. The scale indicates the count numbers of taxa in correlation with the intensity of the shading.

Some taxa were also found to be significantly indicative in these guts (Fig. 6; Tables S6, S7). The majority of indicator genera was found in guts of isopods fed with PLA-food (8 out of 14 and 13 out of 21 genera on 16S rRNA gene and 16S rRNA level, respectively) and none were found in those fed with PS-food. The communities in guts of isopods fed with control-food were more similar to those fed with PLA-food than to those fed with PET- or PS-food, as the former shared more indicator taxa. In particular, most of these genera were more abundant in guts of isopods fed with PLA-food than in those fed with control-food. *Chryseobacterium*, *Devosia*, *Niabella, Prosthecobacter*, *Taeseokella* and uncultured Rhodospirillales were among the indicator taxa on 16S rRNA level in guts of isopods fed with PLA-food (Fig. 6a, Table S7). In guts of isopods fed with PET-food, *Legionella*, *Microbacterium*, *Mycobacterium*, *Paenibacillus* and at least three different genera within the *Enterobacteriaceae* were attributed to indicator genera on 16S rRNA level (Fig. 6b, Table S7).

**Fig. 6:**
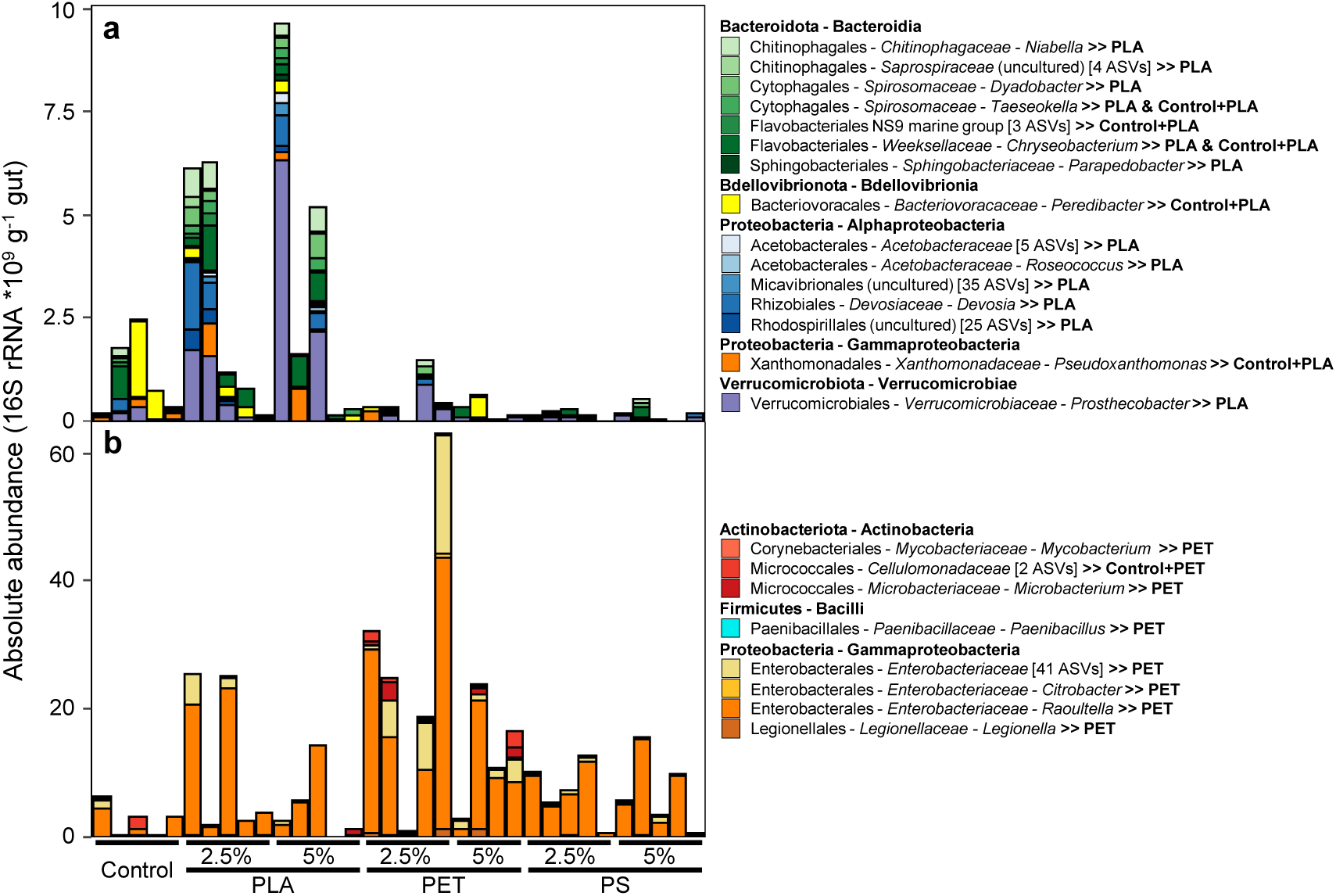
Indicator taxa in isopod guts on 16S rRNA level. The relative abundances of taxa were normalized with the total 16S rRNA abundance derived from qPCR analysis. Indicator taxa for the PLA- (and control-) (**a**) or PET- (and control-) (**b**) treatments were identified. Only taxa on genus level (if applicable, otherwise lowest classification and the number of ASVs are given) that occur in at least three replicates were accepted as potential indicators.

In addition, MP affected food communities. More exclusive genera were obtained in the PS-food than in other kind of food-pellets (Fig. S10b,c). Accordingly, more indicator taxa were found in the PS-food with most of them belonging to the Rhizobiales (Tables S8 and S9). For the control- or PLA-food, no indicators were found. Notably, none of the MP-specific indicator taxa found in the guts were reflected in the respective food-pellets (Tables S7-S10). Such a finding was supported by comparing the abundance of MP-specific indicator taxa from the guts with their abundance in the food pellets (Figs. 6, S11, S12).

## Discussion

Isopods are globally abundant detritivores with a rarely studied gut environment and microbiome, as well as an unknown relevance for atmospheric trace gas emissions. Effects of MP pollution on model detritivores are largely unclear to date. Thus, we provide new insights into effects of biodegradable (PLA) and non-biodegradable (PS and PET) MP polymer types on the gut microbial community as well as activity of the model isopod *P. scaber*, and identify *P. scaber* as a MP-impacted mobile source of molecular hydrogen. We extend previous studies on the effect of PE, tire particles or polyester fibers on life history traits in *P. scaber* that revealed no or only marginal fitness effects [20–22], which is in line with the findings of this study. *P. scaber* was not affected by the ingestion of MP with its food as neither mortality nor weight gain or locomotor activity were altered (Figs. S2, S3). However, when given a choice between control- and MP-food the isopods significantly avoided food containing PS. The palatability of the food source coheres with its microbial composition [14, 15] and therefore it can be considered that PS-food was less attractive, as indeed, most differences of the microbial composition were found between control- and PS-food (Fig. S9). Avoidance behaviour of isopods is commonly observed against metals, pesticides, pharmaceuticals or chars, and already at low concentrations, it is often a more sensitive measure for adverse effects compared to fitness parameters [59–63]. Effects of long-term exposure to MP-contaminated food sources on the isopods’ fitness have not been addressed and cannot be excluded.

Despite the importance of the gut microbiome in soil invertebrates, previous studies testing MP effects on invertebrates are limited [36–39] and are absent in the case of *P. scaber*. In this study, analyses were performed on 16S rRNA gene and 16S rRNA level with the former reflecting the present community and the latter the rather active part of this community, which is commonly a more sensitive response measure. Findings regarding the general gut and food microbiome (MP treatment independent) are discussed in the Supplementary Discussion. The bacterial gut communities of isopods exposed to MP did not differ significantly from the control according to pairwise comparisons (Table S4), but there were differences with respect to exclusive and indicative taxa that were not related to the food communities (Figs. 5, 6, S10-S12, Tables S6-S9). Generally, the gut communities of the isopods fed with MP-free control-food shared more taxa and indicators with those fed with PLA-food than with PS- or PET-food. Moreover, PLA-food increased bacterial proliferation as growth (16S rRNA gene level) and activity (16S rRNA level) were stimulated (Fig. 3). This suggests that some PLA has been degraded in the isopod guts. Abiotic degradation of PLA occurs due to hydrolysis of ester linkages releasing lactic acid at pH 4-7 at low rates [64–66]. The moderately acidic pH inside the gut would thus allow for some abiotic, acid-catalyzed PLA hydrolysis. However, such abiotic hydrolysis takes several months or even years at environmentally relevant temperatures (<30°C) [67]. Biotic PLA degradation is enhanced by enzymatic cleavage of ester bonds and depolymerization of the polymer to oligomers, dimers and lactic acid monomers [68]. Various taxa possess hydrolytic, PLA depolymerizing enzymes like certain lipases, carboxylesterases, and proteinases [69, 70]. Many of such enzymes are active at the gut pH of greater than or equal to 5 [69, 71], suggesting microbial, enzyme-catalyzed rather than abiotic PLA hydrolysis to lactate in the gut of isopods. Subsequent fermentation of lactate in the anoxic gut environment is likely [72, 73].

Indeed, some genera within the Actinobacteria ubiquitous in the isopod guts and less abundant in the food pellets (Fig. S8; Supplementary Discussion) are possible PLA degraders, as numerous members of this class are capable of PLA degradation [74]. Actinobacteria include aerobes as well as anaerobes and may subsist in the mainly anoxic isopod guts [75]. Members of *Alcaligenaceae* were exclusively found in the guts of isopods fed with PLA-food. *Alcaligenes* sp. of the *Alcaligenaceae* are well known to produce lipases that might contribute to PLA hydrolysis [75, 76]. The absence of a massive stimulation of such taxa in PLA treatments in spite of a potential PLA hydrolysis activity might be due to the lack of a specialized metabolism necessary for energy conservation from lactate under anoxic conditions. *Saprospiraceae* (*Parapedobacter*), *Micavibrionaceae* and *Saccharimonadaceae* (TM7) were indicators for guts of isopods exposed to PLA-food (Table S6). *Parapedobacter luteus* of the *Saprospiraceae* is capable of Tween 80 hydrolysis, which has structural similarities to PLA [77]. *Micavibrionaceae* showed affinity for PLA-blended PBAT films [78], suggesting a possible role of both taxa in PLA hydrolysis. *Saccharimonadaceae* (also known as Saccharibacteria, TM7 or clade G6) show an anaerobic lifestyle, were enriched in soil with the structurally related polymer PBAT, are proposed to scavenge small molecular weight carbon during hydrocarbon degradation and host lactate dehydrogenases, suggesting their involvement in lactic acid removal during PLA degradation [79–82]. Additionally, *Xanthomonadaceae* were found in an anaerobic sludge incubation supplemented with PLA [83], suggesting that *Pseudoxanthomonas,* an active genus indicative for guts of PLA-exposed isopods, may have contributed to PLA degradation (Fig. 6). Further most abundant indicators here were *Chryseobacterium*, *Devosia*, *Niabella*, *Prosthecobacter*, *Taeseokella* and uncultured Rhodospirillales. Potential PLA degradation capabilities of these taxa is currently unknown, but cannot be excluded and they may also possess enzymes capable of cleaving the ester bonds of PLA. However, they may have also taken advantage from enhanced lactic acid release during PLA degradation. A facultative lifestyle is conceivable for all of them [84–89].

A further novelty of this study was the assessment of *in situ* hydrogen production in the isopod guts via microsensor measurements. So far such hydrogen production has only been shown for two other soil dwelling invertebrates, the earthworm *Lumbricus terrestris* and the termite *Reticulitermes flavipes* [34, 35]. Hydrogen production in the guts of isopods exposed to PLA-food was higher than in the other guts (Figs. 1, S6, Table S1), suggesting higher fermentative activities in presence of PLA. Degradation of PLA generates lactate (see above), which can then be fermented to acetate, propionate, carbon dioxide and water, or hydrogen [72, 73]. Obligate anaerobes like *Clostridiaceae* are capable of fermenting acetate and propionate to hydrogen and carbon dioxide, but were absent from isopod guts (Fig. S8). Moreover, other hydrogen-producing fermenting gut bacteria, that are often obligate anaerobes [50], were not identified by the sequencing analyses (Fig. S8) and [Fe-Fe]-hydrogenase genes (in the genomic repertoire of these organisms) were not PCR-amplifiable. The absence of obligate anaerobes in the gut system is somewhat surprising compared to other anoxic gut systems [34, 35, 90, 91] and probably owed to the short gut passage time of ∼5 h not allowing for the proliferation of such organisms from ingested inactive forms [92]. However, the short gut passage time will suffice for the activation of fermentation by facultatives. *Enterobacteriacea* are facultatives, well known to produce hydrogen via mixed acid fermentation. An explanation for the stimulated hydrogen production in PLA-fed isopod guts is thus a variation of the mixed acid fermentation pathway by *Enterobacteriaceae* generating ethanol, acetate and formate, and associated formate hydrogen lyase (FHL) catalyzed hydrogen production (Fig. 7) [93]. A ‘proof of principle’ experiment using *Escherichia coli* as a model organism of the *Enterobacteriaceae* has indeed revealed that hydrogen was generated from lactate (see Supplementary Information for Material and Methods and Results; Fig. S13). A pH of 5 to 6 in the median part of the gut, where hydrogen production was high (Figs. 1, S5, S6), represents favourable conditions for FHL activity and formate transformation [93]. *Enterobacteriacea* were ubiquitous as well as active in *P. scaber* guts as was hydrogen production. Energy conservation via the variation of the mixed acid fermentation pathway is only little not leading to detectable stimulation in growth or activity and may explain why *Enterobacteriacea* were not identified as indicators for the PLA-fed isopod guts.

**Fig. 7:**
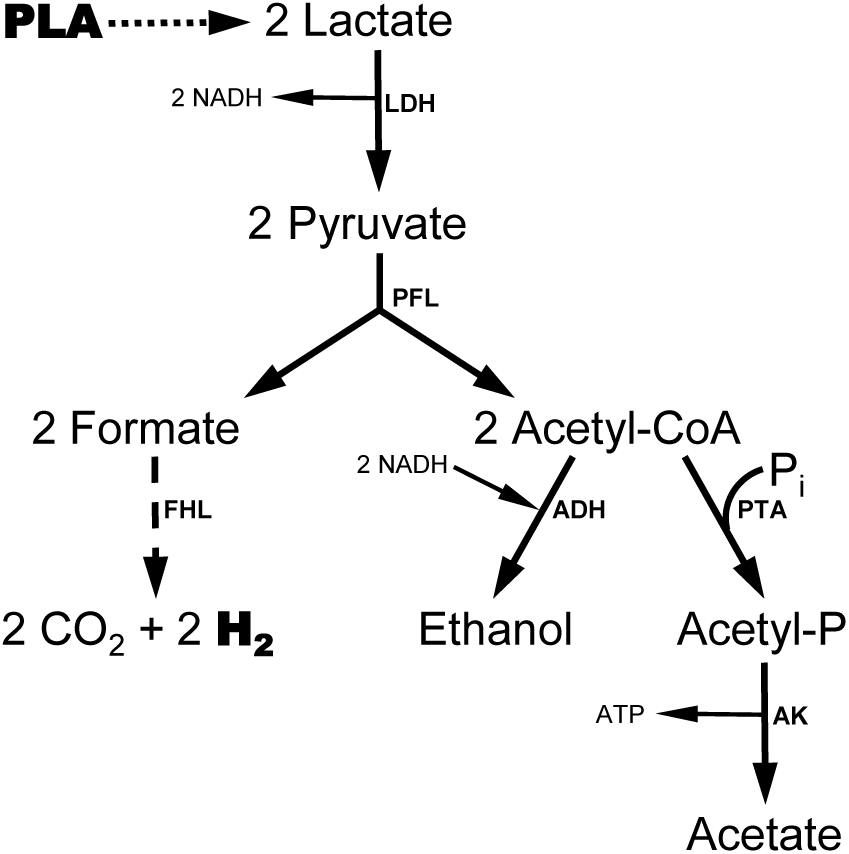
Potential mixed acid fermentation pathway by *Enterobacteriaceae* initiated with lactate. Under conditions, at which pyruvate generated from glucose (or similar compounds) is limited and lactate (that could be derived from PLA as indicated by a dotted arrow) is available, two lactate are oxidized by lactate dehydrogenase (LDH) yielding two pyruvate and two NADH. Two pyruvate react to two acetyl-CoA and two formate catalysed by pyruvate formate lyase (PFL). One acetyl-CoA is then converted to acetate via phosphotransacetylase (PTA) and acetate kinase (AK) yielding one ATP, the other acetyl-CoA is reduced to ethanol by alcohol dehydrogenase (ADH) thereby using two NADH. Formate hydrogen lyase (FHL) transforms formate to carbon dioxide and molecular hydrogen preferentially under acidic conditions (indicated by a dashed arrow).

In contrast, little hydrogen was produced in the guts of isopods fed with PET- and PS-food. Hydrogen consumption by obligate anaerobic methanogens was unlikely to be the reason, as amplification targeting archaeal 16S rRNA failed and methane was not produced (data not shown), when whole isopods were analysed. However, two reasons are conceivable: Either the MP had inhibitory effects on fermentative microorganisms, or Knallgas bacteria were stimulated and consumed most of the hydrogen. Some evidence is given for the latter: *Mycobacterium*, an indicator for the gut of isopods fed with PET-food (Table S8), has been identified as a hydrogen-oxidising bacterium with oxygen as electron acceptor [94]. Oxygen diffusing in the isopod guts from the gut wall might enable hydrogen consumption leading to immediate consumption like well known from termites [41]. Nevertheless, the actual reason of reduced hydrogen emission remains to be determined.

Hydrogen is a valuable electron donor fueling hydrogen-oxidizing processes either inside or outside the gut and MP contamination may have consequences for microbial food webs and global hydrogen emissions. *P. scaber* hydrogen concentrations in the center of the gut ranged from 5 to 30 µM and were thus in the range of those from *L. terrestris* and *R. flavipes*, demonstrating *P. scaber*’s high hydrogen emission potential. Hydrogen emissions from whole isopods were variable, on average 0.83 ± 0.51 ng isopod^-1^ h^-1^ (Fig. 2). Assuming that 20% of the Earth’s terrestrial ecosystems (total surface area of 1.5 x 10^14^ m^2^ [95]) are colonized by these cosmopolitan isopods with a density of 75 isopods m^-2^ (median of distributions given in Paoletti and Hassall [12]), the annual contribution of *P. scabe*r to the global hydrogen production is approximately 0.6 to 2.6 x 10^7^ kg yr^-1^. This value is in the same range of what is annually emitted from paddy fields (1.3 x 10^7^ kg yr^-1^) [96].

Taken together, this study provides new insights regarding the effects of MP on soil invertebrates that are potentially affected by MP-ingestion in the longer term and highlights the hitherto unknown hydrogen emitting capacity of a widely distributed group of detritivores. We identified the moderately acidic, anoxic, median and posterior parts of the isopod’s gut as ‘hot spots’ for hydrogen production. Such a hydrogen production was stimulated by PLA- and inhibited by PET- and PS-ingestion, which was concomitant to changes in the composition of the gut microbiome and in agreement with our initial hypothesis that biodegradable and non-biodegradable MP have contrasting effects. The nature of low hydrogen emissions in response to PET and PS exposure remains speculative, as are consequences of altered hydrogen metabolism inside and outside the isopod gut, opening up new avenues for future research.

## Acknowledgements

This work was funded by the Deutsche Forschungsgemeinschaft (DFG, German Research Foundation); Project Number 391977956; SFB 1357 Microplastic subproject A02. We thank Peter Strohriegl and Lisa Weber for processing polymer granules, Lars Borregard Pedersen for microsensor construction and assistance during measurements, and the Poul Due Jensen Foundation for funding the sensor work. Alina Bernstein and Sabrina Kaupp helped to perform the isopod feeding experiments. We are also grateful to Anja Poehlein for library preparation and sequencing.

## Data availability

Amplicon sequencing data have been deposited in the NCBI Sequence Read Archive (https://www.ncbi.nlm.nih.gov/sra) under the Bioproject PRJNA832915.

