## Supplementary Information for "Effects of microplastic ingestion on hydrogen production and microbiomes in the gut of the terrestrial isopod *Porcellio scaber*"

This document includes:

Supplementary Material and Methods

Supplementary Results

Supplementary Discussion

Supplementary Tables S1 to S10

Supplementary Figures S1 to S15

Supplementary References

**Supplementary Material and Methods**

*Binary food choice experiment*

To evaluate the food choice of the isopods, two food pellets were placed at opposite peripheries of a petri dish with a moist filter paper on the bottom. One of these pellets contained food to be tested (5% MP (PLA, PET or PS) or no MP), while the other contained no MP (control pellet). Before the start of the experiment, all woodlice were starved for three days in boxes containing moist filter paper. Then groups of six woodlice each were placed into petri dishes by releasing the animals one after the other close to the centre of the petri dish. Five groups per combination (no MP vs. no MP; no MP vs. PLA; no MP vs. PET; no MP vs. PS) were monitored with a webcam (Logitech C920, Lausanne, Switzerland) for 48 hrs. Once per hour the number of isopods that were feeding at either food pellet was recorded. The choice index was calculated by subtracting the number of isopods feeding on the control pellets from the number of isopods found at the pellets to be tested, and was divided by the total number of observation where isopods were observed feeding. A negative index represented a preference for the control food, an index around zero no preference and a positive index a preference for the food pellet that was tested.

*Locomotor activity tests*

Non-gravid isopods derived from three randomly chosen jars out of the five jars per treatment were subjected to locomotor activity tests as a measure of their physiological status after two, four and six weeks of the feeding experiment. The isopods were placed on a 30 cm racetrack similar to that described in [1] with four marks every 5 cm (Fig. S14). All runs were carried out in the climate chamber under the same climatic conditions to which they were exposed throughout the experiment. A webcam (Logitech C920, Lausanne, Switzerland) with a frame rate of 15 frames per second was attached over the racetrack to record the runs of the isopods. Each isopod went through three runs with a break of 30 seconds. The videos were evaluated using the software VirtualDub (v1.10.4; www.virtualdub.org), whereby the frames were counted for every 5 cm run until the isopods completely crossed the 5 cm mark. Only the shortest time that was required for an isopod to pass a 5-cm-distance was taken into account for data analysis. For each woodlouse, the body index (weight divided by size) was determined and the speed index was calculated by dividing velocity by body index.

*Sequence analysis*

After demultiplexing of the sequencing reads, bioinformatics were conducted using the QIIME2 software package (version 2021.2; www.qiime2.org) as described previously [2]. Briefly, primer specific sequences were removed from sequencing reads, that were subsequently subjected to the DADA2 pipeline for quality filtering, denoising and joining of paired reads. Low-abundant amplicon sequencing variants (ASVs) with frequencies below the median frequency per ASV of 7 were filtered out and the remaining ASVs were taxonomically classified using a trained classifier based on the SILVA database (version 138; www.arb-silva.de). ASVs shorter than 390 bp and those matching “chloroplast” or “mitochondria” were removed from the data set. Entirely unclassified ASVs or those assigned to unclassified bacteria were manually inspected for chimera formation via NCBI blastn search (www.blast.ncbi.nlm.nih.gov) and excluded in case of the first and the second half of the respective reads was attributed to distanced taxa. If not stated otherwise, ASVs classified as the widespread endosymbionts *Wolbachia* or *Candidatus* Hepatincola [3] likely derived from the gut tissue were also eliminated. The abundance of 16S rRNA gens and 16S rRNA derived from qPCR analyses were re-calculated excluding the proportion of these endosymbionts. With the exception of one sample with less than 5000 reads that was discarded (16S rRNA derived from food containing 2.5% PS), all samples contained at least 20000 reads and further analyzed. Based on the Aitchison distance (a compositional distance metric calculated with the DEICODE plugin [4], the beta diversity of the whole data or of a subset (only food samples, only gut samples) was analyzed. Visualization of the microbial community data was performed with R (version 4.1.1) [5] using the packages *phyloseq* [6] and *ggplot2* [7]. Overlapping (shared) or exclusive (unique) taxa (on genus level, if applicable) in multiple (sub)data sets was demonstrated using Venn diagrams with the r package *ggVennDiagram* [8]. Only taxa that occurred in at least 30% of the respective replicates were included. An indicator species analysis on genus level (if applicable) with the multipatt function in the *indicspecies* package of R (1000 permutations) was conducted to assess specific associations of taxa with specific origin (isopod gut or food) using the relative of 16S rRNA gene and 16S rRNA abundances [9]. The analyses of the associations of taxa with a particular MP-treatment was performed on absolute 16S rRNA gene and 16S rRNA abundances applying the results of the total quantification (qPCR data).

*Test of hydrogen formation from lactate in* Escherichia coli *cultures*

Facultative *Enterobacteriaceae* are frequently found in gut systems, and are well known for their capability to produce molecular hydrogen via mixed acid fermentation.The postulated stimulation of hydrogen formation by *Enterobacteriaceae* via the fermentation of lactate derived from hydrolysis of PLA (see Discussion in the main text) was tested via a pure culture experiment using *Escherichia coli* XL10-Blue. A late exponential culture grown anaerobically at 37°C in M9 minimal medium ([10]; pH 7.2) with 0.4% (w/v) glucose and 0.1% (w/v) yeast extract was used. The culture was washed with glucose-free anoxic M9 medium. Molecular nitrogen stream was applied to minimize contamination with oxygen during decantation and processing. Sealed, airtight 12-mL Exetainer (Labco, Lampeter, UK) containing 8 ml anoxic (flushed with nitrogen) glucose-free M9 medium with 0.01% (w/v) yeast extract and with or without 20 mM lactate added was inoculated resulting in an initial optical density (OD) of ~0.3. About 500 mbar overpressure was applied by injection of 2.5 ml nitrogen gas. Each treatment (with and without lactate) was set up in triplicates. In addition, triplicate sterile medium controls with lactate and triplicate positive controls with additional 0.4% (w/v) glucose were employed. The cultures were incubated at 37°C on a horizontal shaker. Headspace hydrogen concentrations were determined as described above immediately after inoculation, and after ~14 h and ~24 h of incubation. Samples for pH measurements were taken immediately after the gas measurements. OD was measured at 600 nm frequently during the initial 4 h and the second half of the incubation.

### Supplementary Results

*Impact of microplastic on the bacterial communities in the food*

In the food, predominant taxa were attributed to Alpha-, Gammaproteobacteria and Verrucomicrobiae, of which the *Xanthobacteraceae*, *Enterobacteriaceae* and *Opitutaceae* were also highly abundant on 16S rRNA level (Fig. S8e,f). ASVs assigned to *Pseudoxanthomonas spadix, Pseudomonas mohnii* and species within the genus *Novosphingobium* and not further classifiable *Opitutaceae* revealed highest correlations with food rather than gut communities (Fig. S9a). Separation of the communities derived from the different MP-supplemented food pellets was noticeable in the PCoA plot (Fig. S9b). In addition to *Pseudomonas mohnii* and *Opitutaceae*, that were important when analyzing the whole data set (Fig. S9a), ASVs assigned to *Caulobacter, Flavobacterium rivuli* and *Dyadobacter* were highly correlated with differences among food communities (Fig. S9b). Communities on 16S rRNA gene level differed from communities on 16S rRNA level and additionally an interaction effect of MP-treatment and dosage was obtained (PERMANOVA). Investigation of this interaction indicated that most communities differed from each other, with only very few exceptions (Table S5).

*Shared, unique and indicator taxa of food and gut communities regardless the MP-treatment*

Analysis of shared and unique numbers of taxa indicated that 42% of the genera were found in all communities, while 36% were exclusively found in the gut communities with 11% and 5% only found on 16S rRNA gene and 16S rRNA level, respectively (Fig. S10, S15). The indicator analysis confirmed 22 genera as indicators in the isopods guts on both, 16S rRNA gene and 16S rRNA level (Table S10). Another 56 and 5 genera were attributed to indicator taxa on 16S rRNA gene and 16S rRNA level, respectively. Detected indicator taxa in the food-pellets differed from those in the guts. A total of 58 genera were indicative for the food microbiome (16S rRNA gene and/or 16S rRNA level). Noticeably, more genera within the Actinobacteria, Bacteroidia (mostly Chitinophagales and Flavobacteriales), Bacilli, Patescibacteria and

Verrucomicrobiae were indicative in isopod guts than in the food pellets, while more genera within the Acetobacterales, Burkholderiales, Caulobacterales and Enterobacterales were indicative in the food pellets than in the isopod guts.

*Hydrogen formation from lactate in* Escherichia coli

Testing the hydrogen formation by *E. coli* in presence of lactate rather than glucose was performed as a proof of principle. The positive control medium contained glucose and allowed growth without obvious lag-phase (data not shown) indicating that the strain was not impaired due to the harvesting and washing procedure of the pre-culture. Neither growth nor hydrogen production appeared in the sterile negative control (data not shown). In glucose-free cultures with and without lactate added, the OD decreased indicating cell lysis over time (data not shown). Some hydrogen was emitted from cultures incubated without lactate/glucose likely due to the usage of yeast extract or lysed cell components serving as electron donors for living cells. However, more hydrogen was produced in glucose-free cultures in presence of 20 mM lactate. Hydrogen production that was solely attributed to the conversion of lactate was linear over time with a rate of 0.27 nM $h^{-1}$ (Fig. S13). Hence, *E. coli* was indeed capable of hydrogen formation from lactate under anoxic conditions (variation of the mixed acid fermentation), although the rates were low and the reaction appeared not to be growth supportive.

**Supplementary Discussion**

Along with the effects of MP, this study also elucidated the general gut microbiome in comparison with the food microbiome. The 16S rRNA gene abundance revealed that the guts were highly colonized with bacteria, but nearly one order of magnitude less abundant than in the food pellets (Fig. 3). In contrast, Hassall *et al*. [11] demonstrated that up to two orders of magnitude more viable bacterial cells can be obtained from isopods' hindgut content compared to leaves, as was found for soil and gut content for other soil invertebrates like earthworms [12]. Thus, certain food bacteria that are ingested into the gut appear to be selectively digested, but survivors are more prone to proliferate in culture media. Such a conclusion is in agreement with the study of Zimmer and Topp [13] suggesting that a major part of the food microbial community was likely readily digested upon ingestion in the anterior section of the gut. An acidic pH in the anterior section of the gut favours hydrolytic digestive processes, while less acidic pH towards the posterior section of the gut is more suitable for microbial proliferation [14], which agrees with the measured pH values in this study (Fig. S5). It can be speculated that bacteria serving as food source were amongst those that were more abundant in the food than in the gut microbiome, e.g. *Pseudomonas*, *Pseudoxanthomonas* and *Novosphingobium* (Fig. S9a, Table S10), adding such Gram-negative taxa to the list of Gram-positive taxa suggested previously as preferred food [15].

The gut microbiome consisting of mainly Gammaproteobacteria, Bacteroida and Actinobacteria was highly variable (Fig. S8), similar to previous findings [16]. A closer association with the gut microbiome than with the food source appeared for some genera especially within the Actinobacteria, that have been proposed to be important for the immunity of the host [16]. The relative abundance of Verrucomicrobiae was generally higher than in previous analyses [16, 17]. In this study, ~50% of all genera were detected in gut and food communities and amongst others, the Verrucomicrobiae were likely acquired from the food source. However, differences in the beta-diversity of the different food sources were not reflected in the gut communities (Fig. 4). This suggests selective digestion of food taxa and that conditions in the gut restrict the bacterial proliferation to a certain community. Consequently, the gut community possesses higher activities in the gut than in the food, which is in line with the findings regarding the bacterial 16S rRNA abundances in this study (Fig. 3b,e).

### Supplementary Tables

**Table S1**: Minimum pH in the gut of woodlice fed with control food pellets or food pellets containing 5% MP.

| Position | Minimum pH* |
|---|---|
| Anterior | 5.2 (0.5)$^{a}$ |
| Median | 5.8 (0.5)$^{b}$ |
| Posterior | 5.8 (0.6)$^{b}$ |

* Mean values and standard deviations in brackets are shown. Different superscripted letters indicate significant differences in means.

**Table S2**: Maximum hydrogen concentrations in the gut of woodlice fed with control food pellets or food pellets containing 5% MP.

| Position | Maximum hydrogen concentration (µM)* | | | |
|---|---|---|---|---|
| | Control | 5% PLA | 5% PET | 5% PS |
| Anterior | 4.6 (1.6)$^{ab}$ | 6.7 (3.4)$^{ab}$ | 3.4 (1.0)$^{ab}$ | 4.0 (1.0)$^{ab}$ |
| Median | 7.9 (1.8)$^{bc}$ | 30.5 (3.7)$^{d}$ | 2.6 (0.9)$^{a}$ | 5.0 (0.7)$^{ab}$ |
| Posterior | 17.2 (3.8)$^{cd}$ | 24.0 (11.6)$^{d}$ | 5.1 (2.3)$^{ab}$ | 4.3 (2.7)$^{ab}$ |

* Mean values and standard deviations in brackets are shown. Different superscripted letters indicate significant differences in means.

**Table S3**: False discovery rate corrected p-values (1000 permutations) of the pairwise comparisons of the bacterial community composition (based on 16S rRNA gene and 16S rRNA sequences) in the gut of isopods fed with control food pellets or food pellets containing 2.5% or 5% MP. Significant differences ($p < 0.05$) are indicated in bold.

| | Control | 2.5% MP |
|---|---|---|
| 2.5% MP | 0.248 | |
| 5% MP | 0.085 | **0.018** |

**Table S4**: False discovery rate corrected p-values (1000 permutations) of the pairwise comparisons of the bacterial community composition (based on 16S rRNA gene and 16S rRNA sequences) in the gut of isopods fed with control food pellets or food pellets containing PLA, PET or PS.

| | Control | PLA | PET |
|---|---|---|---|
| PLA | 0.337 | | |
| PET | 0.168 | 0.066 | |
| PS | 0.146 | 0.069 | 0.594 |

**Table S5**: False discovery rate corrected p-values (1000 permutations) of the pairwise comparisons of the bacterial community composition (based on 16S rRNA gene and 16S rRNA sequences) in the control food pellets or food pellets containing 2.5% or 5% PLA, PET or PS. Significant differences ($p < 0.05$) are indicated in bold.

| | | Control | PLA | | PET | | PS |
|---|---|---|---|---|---|---|---|
| | | | 2.5% | 5% | 2.5% | 5% | 2.5% |
| PLA | 2.5% | **0.0012** | | | | | |
| | 5% | **0.0012** | **0.0012** | | | | |
| PET | 2.5% | **0.0012** | **0.0012** | **0.0012** | | | |
| | 5% | **0.0070** | 0.1258 | 0.6963 | **0.0012** | | |
| PS | 2.5% | **0.0012** | **0.0012** | **0.0012** | **0.0012** | **0.0012** | |
| | 5% | **0.0012** | **0.0012** | **0.0012** | **0.0012** | **0.0012** | **0.020** |

**Table S6**: Significant indicator genera and respective families in the isopod guts obtained from abundances of 16S rRNA genes normalised with the respective qPCR data. Only taxa were included that occur in at least three replicates.

| Taxon* | $r_{pb}^{g}$ # | p-value | Mean 16S rRNA gene abundance (16S rRNA genes copies $g^{-1}$ gut)[§] | | | |
|---|---|---|---|---|---|---|
| | | | Control | PLA | PET | PS |
| **Control** | | | | | | |
| *Caulobacteraceae - Asticcaulis* | 0.549 | 0.002 | 3.36E+05 | 0.00E+00 | 8.00E+03 | 0.00E+00 |
| **PLA** | | | | | | |
| *Acetobacteraceae - Roseococcus* | 0.517 | 0.018 | 9.71E+03 | 2.90E+05 | 2.12E+04 | 7.61E+04 |
| *Micavibrionaceae* (uncultured) [6] | 0.446 | 0.029 | 0.00E+00 | 4.43E+05 | 0.00E+00 | 0.00E+00 |
| *Microtrichales* (uncultured) [2] | 0.450 | 0.044 | 4.85E+04 | 1.74E+05 | 2.49E+04 | 3.75E+04 |
| Rhodospirillales (uncultured) [25] | 0.509 | 0.022 | 1.42E+04 | 8.60E+05 | 2.50E+04 | 3.02E+04 |
| *Saccharimonadaceae* TM7a [51] | 0.439 | 0.047 | 3.96E+06 | 1.38E+07 | 4.39E+06 | 9.56E+06 |
| *Saprospiraceae* (uncultured) [4] | 0.425 | 0.035 | 5.48E+04 | 2.46E+05 | 1.56E+04 | 0.00E+00 |
| *Sphingobacteriaceae - Parapedobacter* | 0.464 | 0.034 | 2.38E+04 | 5.19E+05 | 3.79E+04 | 2.03E+04 |
| *Terrimicrobiaceae - Terrimicrobium* | 0.468 | 0.042 | 4.05E+06 | 1.39E+07 | 3.01E+06 | 2.08E+06 |
| **PET** | | | | | | |
| *Enterobacteriaceae - Citrobacter* | 0.464 | 0.024 | 1.39E+04 | 2.81E+04 | 3.79E+05 | 6.27E+04 |
| *Propionibacteriaceae - Cutibacterium* | 0.462 | 0.046 | 4.16E+03 | 0.00E+00 | 2.03E+04 | 0.00E+00 |
| *Sphingomonadaceae - Sphingomonas* | 0.423 | 0.048 | 1.22E+06 | 2.07E+06 | 3.44E+06 | 1.88E+06 |
| *Xanthobacteraceae - Bradyrhizobium* | 0.504 | 0.024 | 3.70E+03 | 0.00E+00 | 5.59E+04 | 1.25E+04 |
| *Xanthobacteraceae - Ancylobacter* | 0.441 | 0.046 | 4.03E+04 | 2.40E+05 | 5.61E+05 | 2.91E+05 |
| **Control+PLA** | | | | | | |
| *Bacteriovoracaceae - Peredibacter* | 0.436 | 0.008 | 2.53E+06 | 1.28E+06 | 3.63E+05 | 6.41E+04 |
| *Ca.* Kaiserbacteria [11] | 0.367 | 0.039 | 2.50E+05 | 4.10E+05 | 2.89E+04 | 1.36E+04 |
| *Ca.* Nomurabacteria [5] | 0.318 | 0.035 | 3.28E+06 | 1.43E+06 | 1.73E+05 | 3.23E+04 |
| *Caulobacteraceae - Brevundimonas* | 0.428 | 0.016 | 5.07E+06 | 2.44E+06 | 4.78E+05 | 2.18E+05 |
| *Chitinophagales* (uncultured) [5] | 0.352 | 0.029 | 4.78E+05 | 1.25E+06 | 3.43E+04 | 3.81E+04 |
| *Methylophilaceae - Methylotenera* | 0.393 | 0.004 | 2.33E+05 | 2.81E+05 | 4.70E+03 | 1.11E+04 |
| Nitriliruptoraceae - *Nitriliruptoraceae* | 0.385 | 0.009 | 9.64E+04 | 4.21E+04 | 0.00E+00 | 1.05E+04 |
| *Saprospiraceae* (uncultured) [4] | 0.315 | 0.038 | 5.48E+04 | 2.46E+05 | 1.56E+04 | 0.00E+00 |
| *Sphingomonadaceae* [6] | 0.367 | 0.032 | 4.30E+05 | 7.30E+05 | 8.84E+04 | 0.00E+00 |
| *Spirosomaceae - Taeseokella* | 0.388 | 0.045 | 3.70E+05 | 4.09E+05 | 4.79E+04 | 3.91E+04 |
| *Weeksellaceae - Chryseobacterium* | 0.413 | 0.007 | 2.42E+06 | 2.22E+06 | 8.11E+04 | 1.63E+05 |
| **Control+PET** | | | | | | |
| *Cellulomonadaceae* [2] | 0.310 | 0.045 | 4.56E+06 | 1.05E+06 | 6.63E+06 | 9.42E+05 |

*number of ASVs in parentheses in case of classification on genus level was not applicable

[#]group-equalized point biserial correlation indices

[§]intensity of the green color correlates linearly with the abundance of each taxon; absent taxa are indicated in white

**Table S7**: Significant indicator genera and respective families in the isopod guts obtained from abundances of 16S rRNA normalised with the respective qPCR data. Only taxa were included that occur in at least three replicates.

| **Taxon*** | **$r^{g}_{pb}$ #** | **p-value** | **Mean 16S rRNA abundance (16S rRNA copies g$^{-1}$ gut)§** | | | |
|---|---|---|---|---|---|---|
| | | | **Control** | **PLA** | **PET** | **PS** |
| **Control** | | | | | | |
| *Bacteriovoracaceae - Peredibacter* | 0.456 | 0.031 | 5.21E+08 | 1.37E+08 | 8.21E+07 | 8.13E+06 |
| **PLA** | | | | | | |
| *Acetobacteraceae - Roseococcus* | 0.471 | 0.027 | 6.25E+05 | 1.85E+07 | 0.00E+00 | 8.25E+05 |
| *Acetobacteraceae* [5] | 0.416 | 0.044 | 4.38E+05 | 3.76E+07 | 0.00E+00 | 0.00E+00 |
| *Chitinophagaceae - Niabella* | 0.501 | 0.025 | 4.52E+07 | 2.31E+08 | 2.57E+07 | 1.25E+07 |
| *Devosiaceae - Devosia* | 0.447 | 0.049 | 6.91E+07 | 3.48E+08 | 3.05E+07 | 1.95E+07 |
| *Micavibrionales* (uncultured) [35] | 0.496 | 0.008 | 9.51E+06 | 5.65E+07 | 9.24E+05 | 2.45E+05 |
| *Rhodospirillales* (uncultured) [25] | 0.510 | 0.020 | 6.25E+05 | 1.07E+08 | 1.23E+06 | 1.29E+06 |
| *Saprospiraceae* (uncultured) [4] | 0.430 | 0.038 | 0.00E+00 | 1.84E+08 | 5.51E+06 | 0.00E+00 |
| *Sphingobacteriaceae - Parapedobacter* | 0.463 | 0.043 | 1.40E+05 | 2.64E+07 | 3.94E+06 | 2.66E+06 |
| *Spirosomaceae - Dyadobacter* | 0.461 | 0.042 | 2.07E+07 | 1.54E+08 | 4.21E+07 | 1.63E+07 |
| *Spirosomaceae - Taeseokella* | 0.571 | 0.008 | 1.12E+07 | 5.82E+07 | 2.25E+06 | 1.83E+06 |
| *Verrucomicrobiaceae - Prosthecobacter* | 0.451 | 0.031 | 1.25E+08 | 1.23E+09 | 1.75E+08 | 6.23E+07 |
| *Weeksellaceae - Chryseobacterium* | 0.471 | 0.030 | 8.87E+07 | 1.89E+08 | 1.55E+07 | 2.17E+07 |
| **PET** | | | | | | |
| *Enterobacteriaceae* [41] | 0.514 | 0.010 | 2.30E+08 | 7.38E+08 | 4.38E+09 | 3.85E+08 |
| *Enterobacteriaceae - Citrobacter* | 0.407 | 0.042 | 8.28E+06 | 1.96E+06 | 1.03E+08 | 3.79E+06 |
| *Enterobacteriaceae - Raoultella* | 0.462 | 0.037 | 1.72E+09 | 7.27E+09 | 1.51E+10 | 6.48E+09 |
| *Legionellaceae - Legionella* | 0.456 | 0.046 | 7.53E+07 | 7.87E+07 | 3.75E+08 | 6.49E+07 |
| *Microbacteriaceae - Microbacterium* | 0.495 | 0.016 | 1.15E+08 | 1.13E+08 | 7.41E+08 | 1.28E+08 |
| *Mycobacteriaceae - Mycobacterium* | 0.461 | 0.038 | 4.42E+06 | 4.49E+07 | 1.06E+08 | 2.99E+07 |
| *Paenibacillaceae - Paenibacillus* | 0.508 | 0.008 | 0.00E+00 | 0.00E+00 | 3.08E+07 | 3.66E+05 |
| **Control+PLA** | | | | | | |
| *Bacteriovoracaceae - Peredibacter* | 0.335 | 0.034 | 5.21E+08 | 1.37E+08 | 8.21E+07 | 8.13E+06 |
| Flavobacteriales NS9 marine group [3] | 0.380 | 0.018 | 2.39E+07 | 6.10E+07 | 0.00E+00 | 1.11E+06 |
| *Spirosomaceae - Taeseokella* | 0.406 | 0.042 | 1.12E+07 | 5.82E+07 | 2.25E+06 | 1.83E+06 |
| *Weeksellaceae - Chryseobacterium* | 0.445 | 0.018 | 8.87E+07 | 1.89E+08 | 1.55E+07 | 2.17E+07 |
| *Xanthomonadaceae - Pseudoxanthomonas* | 0.355 | 0.039 | 8.86E+07 | 1.97E+08 | 2.63E+07 | 3.37E+06 |
| **Control+PET** | | | | | | |
| *Cellulomonadaceae* [2] | 0.457 | 0.001 | 5.03E+08 | 4.72E+07 | 6.97E+08 | 3.64E+07 |

*number of ASVs in parentheses in case of classification on genus level was not applicable

#group-equalized point biserial correlation indices

§intensity of the green color correlates linearly with the abundance of each taxon; absent taxa are indicated in white

**Table S8**: Significant indicator genera and respective families in the food pellets obtained from abundances of 16S rRNA genes normalised with the respective qPCR data. Only taxa were included that occur in at least three replicates.

| **Taxon*** | **$r^{g}_{pb}$ #** | **p-value** | **Mean 16S rRNA gene abundance (16S rRNA genes copies $g^{-1}$ food)§** | | | |
|---|---|---|---|---|---|---|
| | | | **Control** | **PLA** | **PET** | **PS** |
| **PET** | | | | | | |
| *Opitutaceae - Lacunisphaera* | 0.451 | 0.024 | 2.09E+04 | 0.00E+00 | 1.65E+06 | 0.00E+00 |
| *Rhodobacteraceae - Gemmobacter* | 0.697 | 0.001 | 5.26E+05 | 9.52E+05 | 2.83E+07 | 3.03E+06 |
| *Rhodobacteraceae - Defluviimonas* | 0.466 | 0.020 | 0.00E+00 | 0.00E+00 | 3.11E+07 | 0.00E+00 |
| *Solimonadaceae* (uncultured) [3] | 0.636 | 0.003 | 0.00E+00 | 0.00E+00 | 3.39E+05 | 0.00E+00 |
| *Spirosomaceae - Leadbetterella* | 0.460 | 0.009 | 0.00E+00 | 0.00E+00 | 6.50E+05 | 1.62E+04 |
| **PS** | | | | | | |
| *Acetobacteraceae - Roseomonas* | 0.448 | 0.042 | 7.80E+06 | 8.85E+06 | 8.77E+06 | 1.90E+07 |
| *Azospirillaceae - Azospirillum* | 0.422 | 0.028 | 0.00E+00 | 1.74E+04 | 1.66E+04 | 2.73E+06 |
| Bacteroidetes VC2.1 Bac22 [3] | 0.426 | 0.026 | 3.80E+05 | 1.19E+05 | 6.14E+03 | 3.15E+06 |
| *Devosiaceae - Devosia* | 0.442 | 0.034 | 8.79E+06 | 1.71E+07 | 1.40E+07 | 2.60E+07 |
| *Kaistiaceae - Kaistia* | 0.646 | 0.001 | 9.88E+06 | 1.81E+07 | 1.04E+07 | 3.63E+07 |
| *Micrococcaceae - Glutamicibacter* | 0.569 | 0.007 | 0.00E+00 | 2.18E+04 | 1.81E+04 | 2.75E+05 |
| *Nocardioidaceae - Aeromicrobium* | 0.484 | 0.022 | 6.02E+05 | 1.07E+06 | 1.21E+06 | 2.55E+06 |
| *Opitutaceae* (uncultured) [8] | 0.475 | 0.030 | 2.75E+06 | 3.11E+06 | 3.08E+07 | 6.75E+07 |
| *Rhizobiaceae - Ochrobactrum* | 0.540 | 0.009 | 1.90E+06 | 2.58E+06 | 4.78E+06 | 1.23E+07 |
| *Solirubrobacteraceae - Patulibacter* | 0.445 | 0.049 | 0.00E+00 | 0.00E+00 | 3.73E+04 | 1.69E+05 |
| *Spirosomaceae - Dyadobacter* | 0.609 | 0.001 | 1.23E+07 | 9.17E+06 | 4.79E+07 | 1.11E+08 |
| *Spirosomaceae - Arcicella* | 0.457 | 0.039 | 2.33E+05 | 3.48E+05 | 1.20E+05 | 1.37E+06 |
| *Weeksellaceae - Chryseobacterium* | 0.482 | 0.015 | 1.75E+06 | 6.27E+06 | 1.48E+07 | 4.15E+07 |
| *Xanthobacteraceae - Ancylobacter* | 0.480 | 0.026 | 3.26E+06 | 2.49E+06 | 3.48E+06 | 7.51E+06 |
| *Xanthobacteraceae - Tardiphaga* | 0.477 | 0.031 | 1.34E+05 | 2.26E+05 | 5.01E+04 | 1.22E+06 |
| *Xanthobacteraceae* [10] | 0.474 | 0.029 | 1.64E+06 | 2.89E+06 | 1.78E+06 | 2.60E+07 |
| **Control+PLA** | | | | | | |
| *Enterobacteriaceae - Raoultella* | 0.494 | 0.006 | 3.99E+08 | 3.55E+08 | 1.43E+08 | 1.22E+08 |
| Oligoflexia 319 6G20 [5] | 0.479 | 0.004 | 1.81E+05 | 1.02E+05 | 0.00E+00 | 0.00E+00 |

*number of ASVs in parentheses in case of classification on genus level was not applicable
#group-equalized point biserial correlation indices
§intensity of the green color correlates linearly with the abundance of each taxon; absent taxa are indicated in white

**Table S9**: Significant indicator genera and respective families in the food pellets obtained from abundances of 16S rRNA normalised with the respective qPCR data. Only taxa were included that occur in at least three replicates.

| **Taxon*** | $r^{g}_{pb}$ # | **p-value** | **Mean 16S rRNA abundance (16S rRNA copies $g^{-1}$ food)§** | | | |
|---|---|---|---|---|---|---|
| | | | **Control** | **PLA** | **PET** | **PS** |
| **PET** | | | | | | |
| *Bacteriovoracaceae - Peredibacter* | 0.468 | 0.030 | 1.06E+05 | 8.91E+05 | 5.93E+06 | 2.41E+05 |
| *Opitutaceae - Lacunisphaera* | 0.442 | 0.024 | 1.42E+04 | 0.00E+00 | 8.14E+06 | 0.00E+00 |
| *Rhodobacteraceae - Gemmobacter* | 0.477 | 0.033 | 8.70E+06 | 1.05E+07 | 1.37E+08 | 7.74E+07 |
| *Yersiniaceae* [1] | 0.461 | 0.033 | 0.00E+00 | 1.76E+04 | 1.38E+06 | 0.00E+00 |
| **PS** | | | | | | |
| *Acetobacteraceae* [5] | 0.481 | 0.030 | 3.26E+07 | 2.01E+07 | 3.52E+07 | 1.11E+08 |
| Bacteroidetes VC2.1 Bac22 [3] | 0.422 | 0.026 | 4.32E+05 | 1.21E+06 | 7.87E+05 | 2.40E+07 |
| *Caulobacteraceae* (uncultured) [11] | 0.549 | 0.011 | 1.25E+07 | 4.61E+07 | 6.91E+07 | 1.56E+08 |
| *Chitinophagaceae - Edaphobaculum* | 0.559 | 0.011 | 3.14E+04 | 1.13E+05 | 1.37E+06 | 6.35E+06 |
| *Devosiaceae - Devosia* | 0.612 | 0.003 | 3.04E+07 | 2.78E+07 | 5.06E+07 | 1.97E+08 |
| *Erwiniaceae* [14] | 0.593 | 0.002 | 9.46E+07 | 1.02E+08 | 2.14E+08 | 1.00E+09 |
| *Kaistiaceae - Kaistia* | 0.596 | 0.006 | 5.95E+07 | 4.46E+07 | 4.54E+07 | 2.41E+08 |
| *Methylopilaceae - Hansschlegelia* | 0.402 | 0.041 | 1.29E+04 | 5.03E+05 | 0.00E+00 | 4.61E+06 |
| *Microbacteriaceae* [40] | 0.530 | 0.014 | 2.85E+07 | 4.84E+07 | 1.54E+08 | 3.37E+08 |
| *Mycobacteriaceae - Mycobacterium* | 0.493 | 0.025 | 3.74E+06 | 4.43E+06 | 9.10E+06 | 1.96E+07 |
| *Pseudomonadaceae - Pseudomonas* | 0.539 | 0.013 | 6.24E+08 | 4.40E+08 | 5.99E+08 | 3.18E+09 |
| *Rhizobiaceae* [14] | 0.534 | 0.012 | 8.81E+06 | 7.61E+06 | 2.05E+07 | 4.32E+07 |
| *Rhizobiaceae - Ochrobactrum* | 0.568 | 0.004 | 9.89E+06 | 5.75E+06 | 8.52E+06 | 1.03E+08 |
| *Rhizobiaceae - Aminobacter* | 0.537 | 0.015 | 7.12E+04 | 5.90E+05 | 9.63E+05 | 3.85E+06 |
| *Rhodobacteraceae - Pseudorhodobacter* | 0.557 | 0.007 | 0.00E+00 | 7.68E+05 | 2.00E+05 | 8.48E+06 |
| *Solirubrobacteraceae - Patulibacter* | 0.678 | 0.001 | 0.00E+00 | 3.12E+05 | 3.19E+05 | 4.30E+06 |
| *Sphingomonadaceae - Sphingomonas* | 0.482 | 0.026 | 1.85E+07 | 1.10E+07 | 1.03E+07 | 5.46E+07 |
| *Spirosomaceae - Arcicella* | 0.494 | 0.023 | 8.04E+05 | 5.83E+06 | 4.53E+06 | 4.57E+07 |
| *Spirosomaceae - Dyadobacter* | 0.466 | 0.039 | 6.11E+07 | 1.72E+07 | 9.51E+07 | 2.50E+08 |
| *Xanthobacteraceae - Ancylobacter* | 0.556 | 0.010 | 1.61E+07 | 1.49E+07 | 1.55E+07 | 9.66E+07 |
| *Xanthobacteraceae - Tardiphaga* | 0.468 | 0.027 | 6.92E+04 | 0.00E+00 | 8.50E+04 | 4.59E+06 |
| *Xanthobacteraceae* [10] | 0.467 | 0.023 | 8.26E+06 | 3.14E+07 | 1.88E+07 | 8.74E+08 |
| *Yersiniaceae - Serratia* | 0.483 | 0.012 | 5.50E+07 | 4.85E+07 | 6.08E+07 | 4.00E+08 |
| **Control+PET** | | | | | | |
| *Acetobacteraceae - Rhodovarius* | 0.419 | 0.030 | 1.06E+07 | 2.33E+06 | 7.63E+06 | 1.15E+06 |

*number of ASVs in parentheses in case of classification on genus level was not applicable

#group-equalized point biserial correlation indices

§intensity of the green color correlates linearly with the abundance of each taxon; absent taxa are indicated in white

**Table S10**: Significant indicator genera and respective families in the isopod guts or the food pellets obtained from the relative abundances of 16S rRNA genes and 16S rRNA. Only taxa were included that occur in at least three replicates.

| **Taxon*** | $r_{pb}^{g}$ # | **p-value** | **Mean relative abundance (%)§** | | | |
|---|---|---|---|---|---|---|
| | | | **Gut** | | **Food** | |
| | | | **16S rRNA gene** | **16S rRNA** | **16S rRNA gene** | **16S rRNA** |
| **Gut - 16S rRNA genes** | | | | | | |
| Absconditabacteriales (SR1) [4] | 0.299 | 0.0041 | 1.70E-03 | 2.47E-04 | 0.00E+00 | 0.00E+00 |
| *Bacillaceae - Anaerobacillus* | 0.574 | 0.0001 | 2.90E-02 | 0.00E+00 | 2.81E-04 | 9.94E-04 |
| *Bacillaceae - Bacillus* | 0.480 | 0.0001 | 9.70E-03 | 2.62E-04 | 0.00E+00 | 1.03E-03 |
| *Beijerinckiaceae - Methylobacterium* | 0.567 | 0.0001 | 5.99E-02 | 1.85E-04 | 1.19E-03 | 2.47E-03 |
| *Beijerinckiaceae* (uncultured) [3] | 0.283 | 0.0133 | 1.22E-03 | 0.00E+00 | 0.00E+00 | 0.00E+00 |
| *Ca.* Campbellbacteria [5] | 0.253 | 0.0001 | 4.46E-02 | 1.99E-03 | 1.79E-04 | 0.00E+00 |
| *Ca.* Kaiserbacteria [11] | 0.213 | 0.0187 | 5.12E-02 | 1.04E-02 | 3.42E-04 | 0.00E+00 |
| *Ca.* Nomurabacteria [5] | 0.246 | 0.0014 | 2.17E-01 | 3.32E-02 | 5.71E-04 | 0.00E+00 |
| *Cellulomonadaceae - Pseudactinotalea* | 0.507 | 0.0001 | 3.81E-02 | 2.33E-03 | 0.00E+00 | 0.00E+00 |
| *Cellvibrionaceae - Cellvibrio* | 0.310 | 0.0009 | 2.61E+00 | 1.19E+00 | 1.18E-01 | 1.07E-01 |
| *Chitinophagaceae - Niabella* | 0.311 | 0.0024 | 3.87E-01 | 1.70E-01 | 3.88E-03 | 7.66E-05 |
| *Chitinophagaceae* - Taibaiella | 0.336 | 0.0001 | 1.25E+00 | 2.59E-01 | 2.84E-01 | 1.51E-02 |
| *Dermacoccaceae* [4] | 0.425 | 0.0001 | 6.75E-02 | 3.89E-03 | 1.43E-04 | 0.00E+00 |
| *Dermacoccaceae - Demetria* | 0.359 | 0.0001 | 5.60E-02 | 1.86E-03 | 0.00E+00 | 0.00E+00 |
| *Enterobacteriaceae - Escherichia* | 0.299 | 0.0022 | 7.53E-03 | 1.90E-03 | 0.00E+00 | 0.00E+00 |
| *Erysipelotrichaceae - Erysipelothrix* | 0.558 | 0.0001 | 2.78E-02 | 0.00E+00 | 3.06E-04 | 1.94E-03 |
| Gaiellales (uncultured) [4] | 0.315 | 0.0001 | 1.39E-01 | 3.11E-02 | 1.62E-03 | 0.00E+00 |
| *Gammaproteobacteria* [7] | 0.337 | 0.0004 | 2.58E-02 | 9.46E-03 | 8.65E-05 | 5.84E-04 |
| *Geodermatophilaceae - Antricoccus* | 0.349 | 0.0003 | 1.14E-02 | 1.31E-03 | 0.00E+00 | 0.00E+00 |
| *Halomonadaceae - Halomonas* | 0.375 | 0.0001 | 4.47E-02 | 1.26E-02 | 8.79E-05 | 5.14E-04 |
| *Ilumatobacteraceae* [4] | 0.329 | 0.0007 | 2.23E-02 | 7.44E-03 | 0.00E+00 | 0.00E+00 |
| *Ilumatobacteraceae* (uncultured) [5] | 0.336 | 0.0005 | 3.89E-03 | 2.44E-04 | 0.00E+00 | 0.00E+00 |
| *Intrasporangiaceae - Ornithinimicrobium* | 0.452 | 0.0001 | 5.79E-02 | 5.27E-03 | 0.00E+00 | 0.00E+00 |
| *Methylophilaceae - Methylotenera* | 0.259 | 0.0128 | 2.36E-02 | 9.34E-03 | 0.00E+00 | 0.00E+00 |
| *Microbacteriaceae - Agromyces* | 0.409 | 0.0001 | 1.91E-02 | 8.63E-04 | 3.15E-04 | 0.00E+00 |
| *Microbacteriaceae - Galbitalea* | 0.382 | 0.0001 | 3.29E-01 | 3.64E-02 | 5.73E-03 | 2.67E-03 |
| *Microbacteriaceae - Leucobacter* | 0.246 | 0.0201 | 1.17E-02 | 4.11E-03 | 0.00E+00 | 0.00E+00 |
| *Microbacteriaceae - Schumannella* | 0.384 | 0.0001 | 3.65E-02 | 2.33E-03 | 1.24E-02 | 1.37E-03 |
| *Micrococcaceae - Glutamicibacter* | 0.234 | 0.0240 | 4.63E-02 | 1.84E-02 | 4.93E-03 | 1.97E-03 |
| *Moraxellaceae - Enhydrobacter* | 0.161 | 0.0146 | 1.15E-02 | 0.00E+00 | 0.00E+00 | 1.02E-04 |
| *Mycobacteriaceae - Mycobacterium* | 0.393 | 0.0002 | 2.49E-01 | 8.29E-02 | 6.41E-02 | 1.29E-01 |
| *Nakamurellaceae - Nakamurella* | 0.594 | 0.0001 | 4.79E-02 | 7.52E-03 | 8.07E-04 | 1.68E-04 |
| *Nitriliruptoraceae - Nitriliruptoraceae* | 0.259 | 0.0001 | 6.39E-03 | 4.94E-04 | 0.00E+00 | 0.00E+00 |
| *Nocardiaceae - Gordonia* | 0.364 | 0.0001 | 2.95E-01 | 9.80E-02 | 7.52E-04 | 0.00E+00 |
| *Nocardiaceae - Rhodococcus* | 0.343 | 0.0004 | 1.08E+00 | 4.05E-01 | 2.85E-03 | 3.20E-03 |
| *Nocardiaceae - Williamsia* | 0.421 | 0.0001 | 2.60E-01 | 6.59E-02 | 1.97E-03 | 2.20E-03 |
| *Nocardioidaceae - Aeromicrobium* | 0.690 | 0.0001 | 2.10E-01 | 2.89E-02 | 5.19E-02 | 2.08E-02 |
| *Nocardioidaceae - Nocardioides* | 0.681 | 0.0001 | 9.86E-01 | 9.45E-02 | 6.20E-02 | 1.06E-02 |
| *Propionibacteriaceae - Cutibacterium* | 0.257 | 0.0121 | 1.90E-03 | 0.00E+00 | 0.00E+00 | 0.00E+00 |
| *Rhizobiaceae - Phyllobacterium* | 0.213 | 0.0447 | 2.83E-03 | 8.69E-04 | 0.00E+00 | 0.00E+00 |
| *Rhodobacteraceae - Amaricoccus* | 0.236 | 0.0307 | 1.30E-02 | 5.77E-03 | 3.95E-04 | 6.73E-04 |
| *Rubritaleaceae - Luteolibacter* | 0.504 | 0.0001 | 1.96E+00 | 6.83E-03 | 3.96E-01 | 2.22E-03 |
| *Rubritaleaceae - Roseibacillus* | 0.371 | 0.0001 | 1.08E-02 | 0.00E+00 | 0.00E+00 | 0.00E+00 |
| *Rubritaleaceae - Rubritalea* | 0.288 | 0.0001 | 5.88E-02 | 0.00E+00 | 0.00E+00 | 0.00E+00 |
| *Saccharimonadaceae* [1] | 0.256 | 0.0037 | 2.16E-02 | 0.00E+00 | 0.00E+00 | 0.00E+00 |
| *Saccharimonadaceae* TM7a [51] | 0.710 | 0.0001 | 1.55E+00 | 2.20E-02 | 4.37E-02 | 9.65E-04 |
| Saccharimonadales [7] | 0.635 | 0.0001 | 2.37E+00 | 6.77E-02 | 1.62E-01 | 5.21E-03 |
| Saccharimonadales LWQ8 [3] | 0.474 | 0.0001 | 4.11E-01 | 1.73E-03 | 1.39E-03 | 1.12E-04 |
| *Sphingomonadaceae - Altererythrobacter* | 0.222 | 0.0159 | 5.55E-02 | 6.64E-03 | 1.33E-02 | 2.57E-03 |
| *Sphingomonadaceae - Sphingopyxis* | 0.264 | 0.0119 | 1.33E-02 | 2.19E-03 | 0.00E+00 | 0.00E+00 |
| Thermomicrobiales JG30 KF CM45 [10] | 0.493 | 0.0001 | 1.82E-01 | 1.71E-02 | 8.18E-04 | 2.72E-04 |
| Verrucomicrobiales DEV007 [3] | 0.427 | 0.0001 | 5.72E-02 | 6.71E-04 | 4.01E-03 | 0.00E+00 |
| *Weeksellaceae - Cloacibacterium* | 0.234 | 0.0133 | 3.25E-03 | 0.00E+00 | 0.00E+00 | 4.81E-04 |
| *Xanthobacteraceae - Bradyrhizobium* | 0.296 | 0.0008 | 8.10E-03 | 0.00E+00 | 0.00E+00 | 0.00E+00 |
| *Xanthomonadaceae - Luteimonas* | 0.271 | 0.0083 | 1.39E-01 | 4.77E-02 | 2.36E-04 | 2.99E-04 |
| *Xanthomonadaceae - Stenotrophomonas* | 0.204 | 0.0414 | 1.82E-02 | 1.75E-03 | 1.98E-03 | 2.27E-03 |

**Table S10**: *continued*

| Taxon* | $r^{g}_{pb}$ # | p-value | Mean relative abundance (%)§ | | | |
|---|---|---|---|---|---|---|
| | | | Gut | | Food | |
| | | | 16S rRNA gene | 16S rRNA | 16S rRNA gene | 16S rRNA |
| **Gut - 16S rRNA** | | | | | | |
| Flavobacteriales NS9 marine group [3] | 0.239 | 0.0098 | 9.44E-03 | 6.04E-02 | 0.00E+00 | 0.00E+00 |
| *Flavobacteriaceae - Aurantivirga* | 0.188 | 0.0205 | 9.93E-03 | 9.02E-02 | 0.00E+00 | 1.30E-04 |
| *Flavobacteriaceae - Maribacter* | 0.246 | 0.0240 | 1.42E-02 | 4.78E-02 | 0.00E+00 | 1.79E-04 |
| *Verrucomicrobiaceae - Verrucomicrobium* | 0.248 | 0.0221 | 9.14E-02 | 2.29E-01 | 6.81E-02 | 8.43E-02 |
| *Xanthomonadaceae* [2] | 0.206 | 0.0489 | 6.58E-05 | 1.28E-03 | 0.00E+00 | 0.00E+00 |
| **Gut - 16S rRNA genes+16S rRNA** | | | | | | |
| *Beijerinckiaceae - Bosea* | 0.253 | 0.0155 | 3.30E-02 | 3.00E-02 | 3.66E-03 | 1.78E-02 |
| *Beijerinckiaceae - Camelimonas* | 0.238 | 0.0245 | 7.16E-02 | 6.04E-02 | 1.48E-04 | 2.44E-04 |
| *Cellulomonadaceae* [2] | 0.284 | 0.0042 | 5.66E-01 | 5.13E-01 | 1.32E-02 | 6.40E-03 |
| Chitinophagales (uncultured) [5] | 0.283 | 0.0060 | 6.53E-02 | 4.01E-02 | 0.00E+00 | 0.00E+00 |
| *Comamonadaceae* [82] | 0.303 | 0.0013 | 7.39E-01 | 5.48E-01 | 1.60E-01 | 2.28E-01 |
| *Dermabacteraceae - Brachybacterium* | 0.315 | 0.0014 | 5.78E-02 | 2.86E-02 | 3.30E-04 | 0.00E+00 |
| Entomoplasmatales - *Ca.* Hepatoplasma | 0.361 | 0.0003 | 1.89E+00 | 1.33E+00 | 3.99E-02 | 9.59E-03 |
| *Flavobacteriaceae* [16] | 0.343 | 0.0007 | 9.48E-02 | 1.02E-01 | 0.00E+00 | 0.00E+00 |
| *Flavobacteriaceae - Mesonia* | 0.268 | 0.0118 | 7.14E-03 | 7.58E-03 | 0.00E+00 | 0.00E+00 |
| *Francisellaceae - Francisella* | 0.252 | 0.0163 | 7.25E-03 | 8.04E-03 | 1.01E-04 | 0.00E+00 |
| *Iamiaceae - Iamia* | 0.469 | 0.0001 | 2.45E-02 | 1.66E-02 | 3.51E-04 | 1.46E-04 |
| *Ilumatobacteraceae - Ilumatobacter* | 0.259 | 0.0123 | 4.72E-02 | 2.55E-02 | 1.52E-04 | 0.00E+00 |
| *Microbacteriaceae - Microbacterium* | 0.337 | 0.0006 | 5.29E-01 | 4.46E-01 | 1.48E-01 | 5.40E-02 |
| *Rhodobacteraceae - Paracoccus* | 0.496 | 0.0001 | 1.39E+00 | 1.30E+00 | 4.11E-02 | 9.86E-02 |
| *Saprospiraceae* (uncultured) [4] | 0.247 | 0.0213 | 9.97E-03 | 1.50E-02 | 0.00E+00 | 0.00E+00 |
| *Solirubrobacteraceae - Patulibacter* | 0.268 | 0.0092 | 1.48E-02 | 1.72E-02 | 1.94E-03 | 9.70E-03 |
| *Sphingobacteriaceae - Parapedobacter* | 0.267 | 0.0099 | 2.83E-02 | 1.36E-02 | 0.00E+00 | 0.00E+00 |
| *Spirosomaceae - Taeseokella* | 0.287 | 0.0050 | 4.18E-02 | 3.82E-02 | 1.76E-04 | 0.00E+00 |
| *Terrimicrobiaceae - Terrimicrobium* | 0.385 | 0.0001 | 9.10E-01 | 1.01E+00 | 8.95E-02 | 1.39E-01 |
| *Verrucomicrobiaceae* (uncultured) [10] | 0.300 | 0.0012 | 5.27E-02 | 4.16E-02 | 1.63E-03 | 7.34E-03 |
| *Vibrionaceae - Vibrio* | 0.619 | 0.0001 | 2.27E+01 | 4.60E+01 | 3.31E-02 | 4.06E-01 |
| *Weeksellaceae - Moheibacter* | 0.225 | 0.0385 | 1.15E-01 | 1.81E-01 | 1.18E-03 | 5.80E-04 |
| **Food - 16S rRNA genes** | | | | | | |
| *Alcaligenaceae - Pigmentiphaga* | 0.674 | 0.0001 | 5.83E-03 | 2.65E-03 | 1.58E-01 | 7.07E-02 |
| *Bdellovibrionaceae - Bdellovibrio* | 0.532 | 0.0001 | 6.92E-01 | 7.21E-01 | 4.23E+00 | 7.11E-01 |
| *Caulobacteraceae - Brevundimonas* | 0.346 | 0.0002 | 3.14E-01 | 2.17E-01 | 9.69E-01 | 3.48E-01 |
| *Micavibrionales* (uncultured) [35] | 0.292 | 0.0013 | 1.77E-01 | 2.03E-02 | 4.00E-01 | 1.42E-02 |
| *Microbacteriaceae - Herbiconiux* | 0.305 | 0.0026 | 6.42E-04 | 0.00E+00 | 7.20E-03 | 1.74E-03 |
| *Oxalobacteraceae - Herminiimonas* | 0.406 | 0.0001 | 9.73E-03 | 1.62E-03 | 3.53E-02 | 4.73E-03 |
| *Solimonadaceae* (uncultured) [3] | 0.304 | 0.0025 | 2.92E-04 | 8.53E-04 | 6.67E-03 | 1.52E-03 |
| *Sphingobacteriaceae* [2] | 0.300 | 0.0131 | 0.00E+00 | 0.00E+00 | 6.12E-04 | 0.00E+00 |
| *Sphingobacteriaceae - Pedobacter* | 0.310 | 0.0016 | 5.51E-01 | 8.28E-01 | 2.03E+00 | 1.13E+00 |
| *Spirosomaceae - Dyadobacter* | 0.528 | 0.0001 | 6.87E-02 | 1.08E-01 | 1.66E+00 | 5.85E-01 |
| *Verrucomicrobiae* (uncultured) [5] | 0.495 | 0.0001 | 1.79E-02 | 2.62E-02 | 2.44E-01 | 1.03E-01 |
| *Weeksellaceae - Chryseobacterium* | 0.270 | 0.0070 | 2.30E-01 | 2.17E-01 | 5.77E-01 | 1.62E-01 |
| *Yersiniaceae - Serratia* | 0.624 | 0.0001 | 1.68E-01 | 3.01E-02 | 5.28E+00 | 1.31E+00 |
| **Food - 16S rRNA** | | | | | | |
| *Acetobacteraceae* [5] | 0.648 | 0.0001 | 7.38E-03 | 1.12E-02 | 1.51E-01 | 4.35E-01 |
| *Acetobacteraceae - Rhodovarius* | 0.397 | 0.0001 | 1.18E-02 | 6.83E-03 | 3.37E-02 | 7.80E-02 |
| *Acetobacteraceae - Roseococcus* | 0.457 | 0.0001 | 2.57E-02 | 1.00E-02 | 2.90E-02 | 7.73E-02 |
| *Acetobacteraceae - Roseomonas* | 0.737 | 0.0001 | 1.03E-02 | 6.36E-03 | 4.53E-01 | 1.43E+00 |
| *Acidobacteriaceae - Terriglobus* | 0.261 | 0.0005 | 0.00E+00 | 0.00E+00 | 6.78E-03 | 5.54E-02 |
| *Alphaproteobacteria* [1] | 0.254 | 0.0116 | 0.00E+00 | 0.00E+00 | 1.56E-03 | 4.32E-03 |
| *Bacteriovoracaceae - Bacteriovorax* | 0.345 | 0.0001 | 1.40E-02 | 2.75E-02 | 1.13E-01 | 2.97E-01 |
| *Beijerinckiaceae - Beijerinckiaceae* | 0.448 | 0.0001 | 1.01E-03 | 1.24E-03 | 8.58E-04 | 1.81E-02 |
| *Caulobacteraceae - Caulobacter* | 0.400 | 0.0001 | 1.48E-01 | 1.58E-01 | 1.42E+00 | 4.71E+00 |
| *Comamonadaceae - Aquabacterium* | 0.215 | 0.0120 | 0.00E+00 | 0.00E+00 | 0.00E+00 | 1.66E-03 |
| *Opitutaceae* [6] | 0.254 | 0.0172 | 1.77E-01 | 2.28E-01 | 3.94E-01 | 1.47E+00 |
| *Opitutaceae* (uncultured) [8] | 0.359 | 0.0001 | 8.68E-02 | 1.67E-01 | 8.09E-01 | 3.07E+00 |
| Puniceispirillales (uncultured) [3] | 0.503 | 0.0001 | 8.52E-04 | 1.23E-03 | 3.79E-03 | 2.46E-02 |
| *Rhizobiaceae - Aminobacter* | 0.377 | 0.0002 | 1.65E-03 | 0.00E+00 | 6.23E-03 | 1.40E-02 |
| *Rhodobacteraceae - Defluviimonas* | 0.232 | 0.0349 | 0.00E+00 | 1.10E-02 | 2.22E-01 | 7.11E-01 |
| *Rhodobacteraceae - Pseudorhodobacter* | 0.276 | 0.0058 | 3.90E-03 | 0.00E+00 | 5.05E-03 | 2.42E-02 |
| *Solimonadaceae - Hydrocarboniphaga* | 0.554 | 0.0001 | 4.91E-02 | 3.25E-02 | 8.82E-01 | 2.36E+00 |
| *Xanthobacteraceae* [10] | 0.252 | 0.0210 | 2.81E-01 | 2.97E-01 | 2.75E-01 | 1.02E+00 |
| *Xanthobacteraceae - Azorhizobium* | 0.558 | 0.0001 | 1.01E+00 | 1.15E+00 | 2.97E+00 | 1.19E+01 |

**Table S10**: *continued*

| Taxon* | $r^{g}_{pb}$ # | p-value | Mean relative abundance (%)§ | | | |
|---|---|---|---|---|---|---|
| | | | Gut | | Food | |
| | | | 16S rRNA gene | 16S rRNA | 16S rRNA gene | 16S rRNA |
| **Food - 16S rRNA genes+16S rRNA** | | | | | | |
| *Alcaligenaceae - Verticiella* | 0.426 | 0.0001 | 1.63E-03 | 2.01E-03 | 5.15E-02 | 6.42E-02 |
| *Alteromonadaceae - Rheinheimera* | 0.229 | 0.0286 | 4.39E-03 | 2.40E-03 | 8.36E-02 | 5.21E-02 |
| *Caulobacteraceae - Phenylobacterium* | 0.260 | 0.0130 | 1.03E-02 | 6.88E-03 | 2.96E-02 | 2.32E-02 |
| *Caulobacteraceae* (uncultured) [11] | 0.608 | 0.0001 | 1.28E-02 | 7.56E-03 | 5.46E-01 | 6.75E-01 |
| *Comamonadaceae - Comamonas* | 0.397 | 0.0002 | 5.17E-02 | 1.36E-02 | 2.13E-01 | 2.04E-01 |
| *Comamonadaceae - Variovorax* | 0.737 | 0.0001 | 3.15E-02 | 9.53E-03 | 6.31E-01 | 5.46E-01 |
| *Devosiaceae - Devosia* | 0.459 | 0.0001 | 1.21E-01 | 2.13E-01 | 6.18E-01 | 5.03E-01 |
| Enterobacterales [16] | 0.231 | 0.0339 | 3.12E-02 | 1.88E-02 | 9.80E-02 | 6.71E-02 |
| *Enterobacteriaceae* [41] | 0.351 | 0.0003 | 1.69E+00 | 2.26E+00 | 4.45E+00 | 4.87E+00 |
| *Erwiniaceae* [14] | 0.542 | 0.0001 | 3.21E-02 | 1.06E-03 | 2.64E+00 | 3.02E+00 |
| *Erwiniaceae - Erwinia* | 0.437 | 0.0001 | 9.46E-03 | 0.00E+00 | 1.29E-01 | 1.18E-01 |
| *Methylophilaceae - Methylophilus* | 0.337 | 0.0008 | 7.05E-02 | 7.09E-02 | 3.11E-01 | 2.44E-01 |
| *Oxalobacteraceae* [28] | 0.485 | 0.0001 | 1.17E-01 | 3.62E-02 | 7.26E-01 | 3.85E-01 |
| *Oxalobacteraceae - Duganella* | 0.370 | 0.0001 | 1.94E-02 | 6.51E-03 | 1.24E-01 | 1.04E-01 |
| *Oxalobacteraceae - Herbaspirillum* | 0.554 | 0.0001 | 8.02E-03 | 2.36E-03 | 3.49E-01 | 2.06E-01 |
| Oxalobacteraceae - *Janthinobacterium* | 0.262 | 0.0111 | 3.02E-04 | 5.64E-04 | 1.88E-02 | 1.19E-02 |
| *Pseudomonadaceae - Pseudomonas* | 0.514 | 0.0001 | 1.89E+00 | 1.73E+00 | 1.06E+01 | 8.79E+00 |
| *Rhizobiaceae* [14] | 0.472 | 0.0001 | 3.69E-02 | 2.16E-02 | 1.51E-01 | 1.57E-01 |
| *Rhizobiaceae - Allorhizobium* | 0.632 | 0.0001 | 1.10E+00 | 3.81E-01 | 3.00E+00 | 2.98E+00 |
| *Rhodobacteraceae - Falsirhodobacter* | 0.559 | 0.0001 | 1.55E-02 | 3.25E-03 | 2.91E-01 | 3.48E-01 |
| *Rhodobacteraceae - Ketogulonicigenium* | 0.612 | 0.0001 | 5.00E-02 | 2.04E-03 | 6.51E-01 | 4.24E-01 |
| *Sphingobacteriaceae - Mucilaginibacter* | 0.261 | 0.0099 | 1.97E-02 | 1.24E-02 | 8.51E-02 | 5.06E-02 |
| *Sphingomonadaceae - Novosphingobium* | 0.718 | 0.0001 | 4.78E-01 | 3.39E-02 | 1.59E+01 | 8.24E+00 |
| *Xanthobacteraceae - Ancylobacter* | 0.449 | 0.0001 | 7.36E-02 | 3.07E-02 | 1.69E-01 | 2.39E-01 |
| *Xanthobacteraceae - Tardiphaga* | 0.299 | 0.0024 | 6.05E-04 | 0.00E+00 | 1.46E-02 | 6.98E-03 |
| *Xanthomonadaceae - Pseudoxanthomonas* | 0.629 | 0.0001 | 3.97E-01 | 2.36E-01 | 6.67E+00 | 4.14E+00 |

*number of ASVs in parentheses in case of classification on genus level was not applicable

#group-equalized point biserial correlation indices

§intensity of the green color correlates linearly with the abundance of each taxon; absent taxa are indicated in white

### Supplementary Figures

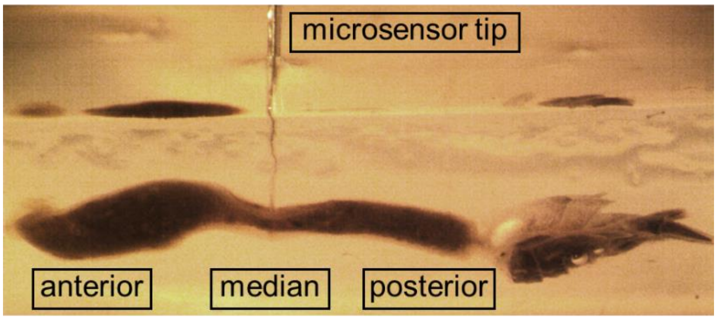


**Fig. S1: Gut of *P. scaber* embedded in agarose.** Anterior, median and posterior positions in the gut are indicated. The picture was taken during a radial microsensor measurement through the median of the gut.

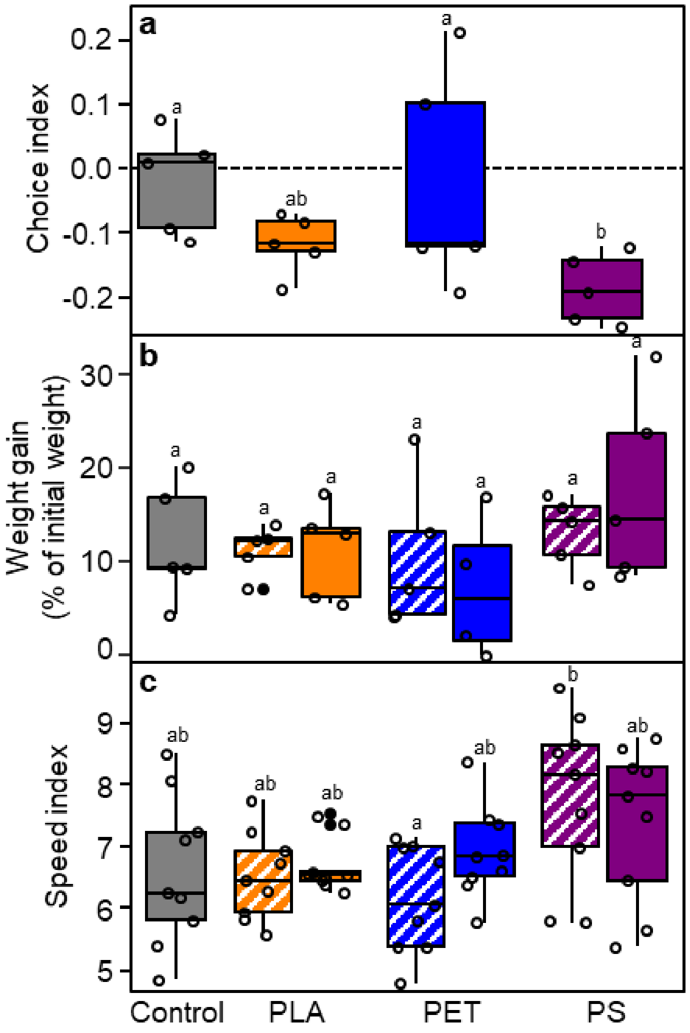


**Fig. S2: Effects of MP ingestion on the avoidance behavior and fitness parameters of *P. scaber.*** Isopods were exposed to control-food pellets or food pellets containing PLA, PET or PS. a presents data derived from a binary food choice experiment performed with 5 isopods each. Negative choice index values indicate preferences for control-food, positive choice index values indicate preferences for 5%-MP-food. b, c present data derived during an eight-week exposure to control-, 2.5%-MP- (striped) or 5%-MP-food (no pattern). The weight gain of isopods in five jars per treatment was assessed after eight weeks (b). After two, four and six weeks average speed indexes of non-gravid isopods derived from three different jars per treatment were determined (c). Open circles indicate data points, closed black circles indicate outliers and different lower case letters above boxplots indicate significant differences in means (a, b, c; $p < 0.05$).

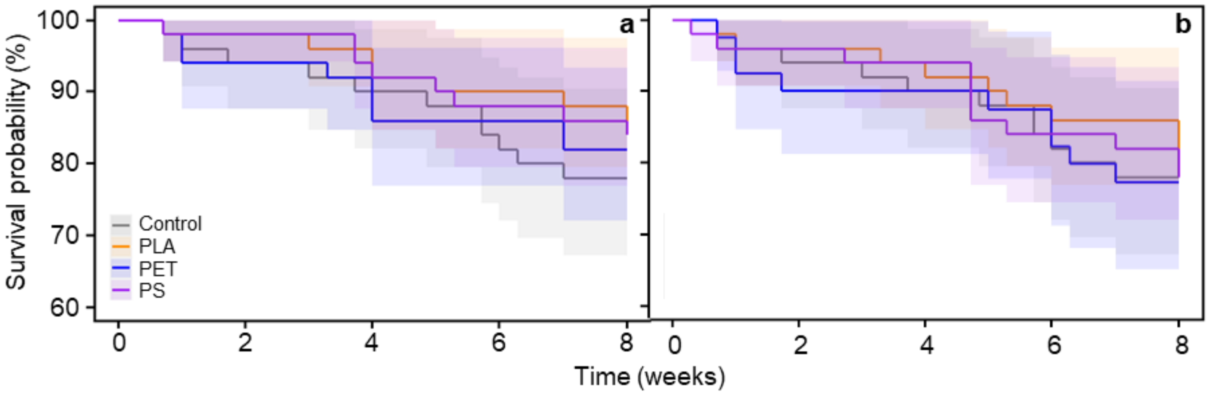


**Fig. S3: Effects of MP ingestion on the survival of *P. scaber.*** Isopods were exposed to control-food pellets or food pellets containing 2.5% (a) or 5% (b) PLA, PET or PS. Survival was examined at least every other day during an eight-week feeding experiment. Step functions and associated 95% confidence intervals of isopods derived from five different jars per treatment are plotted. Overlapping confidence intervals indicate no significant differences.

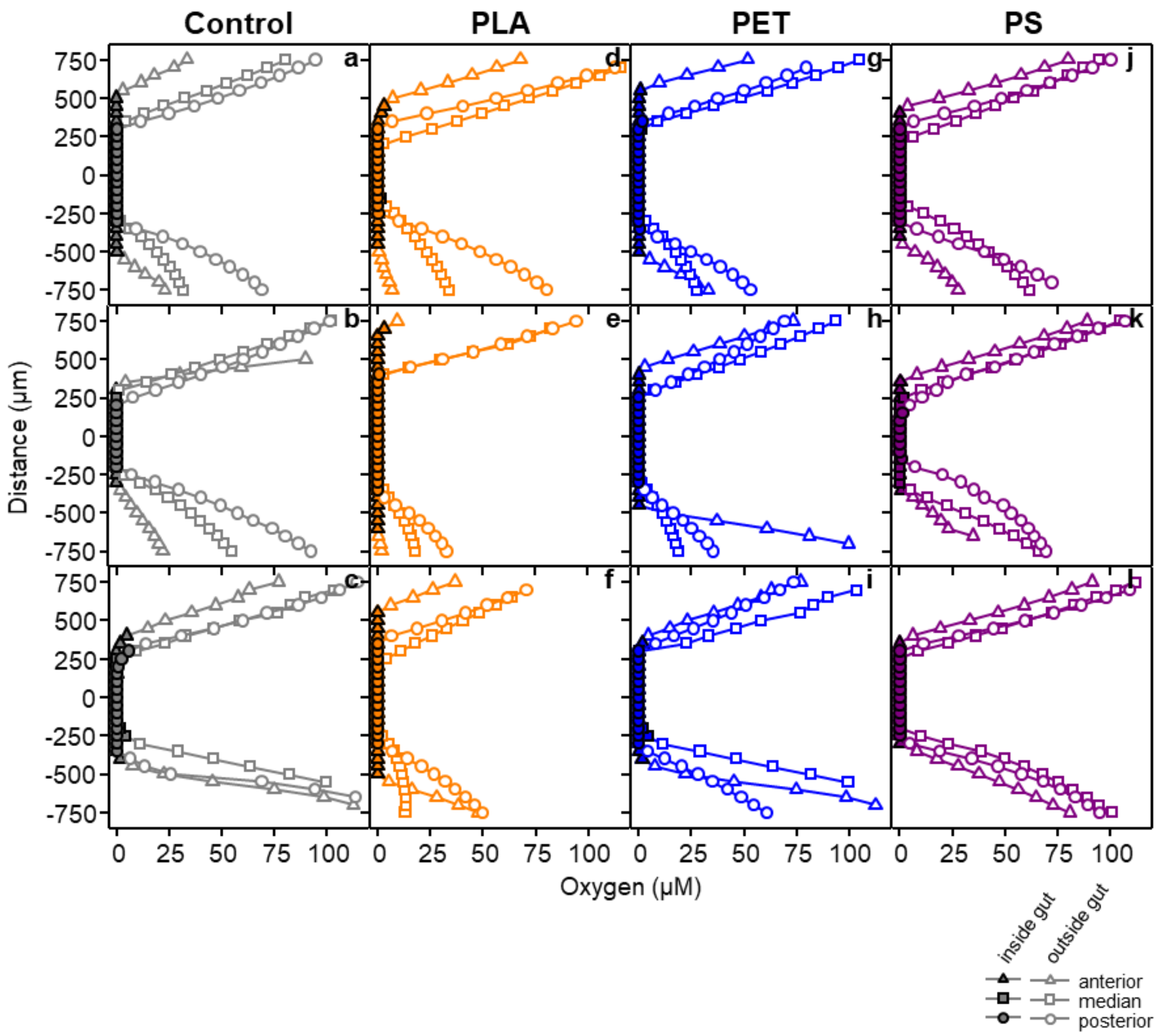


**Fig. S4: Radial oxygen profiles of woodlice guts.** The woodlice were fed with food containing no microplastic particles (control; **a**, **b**, **c**) or 5% PLA (**d**, **e**, **f**), PET (**g**, **h**, **i**) or PS (**j**, **k**, **l**) for six days prior gut extraction and subsequent embedding in agarose and microsensor measurements. For each gut, profiles were recorded from the anterior, median and posterior. Closed and open symbols represent measured concentrations inside and outside (in agarose) the guts, respectively. Each panel with the same color represents one of three replicate guts per treatment. The distance of 0 µm indicates the center of the tube-like gut.

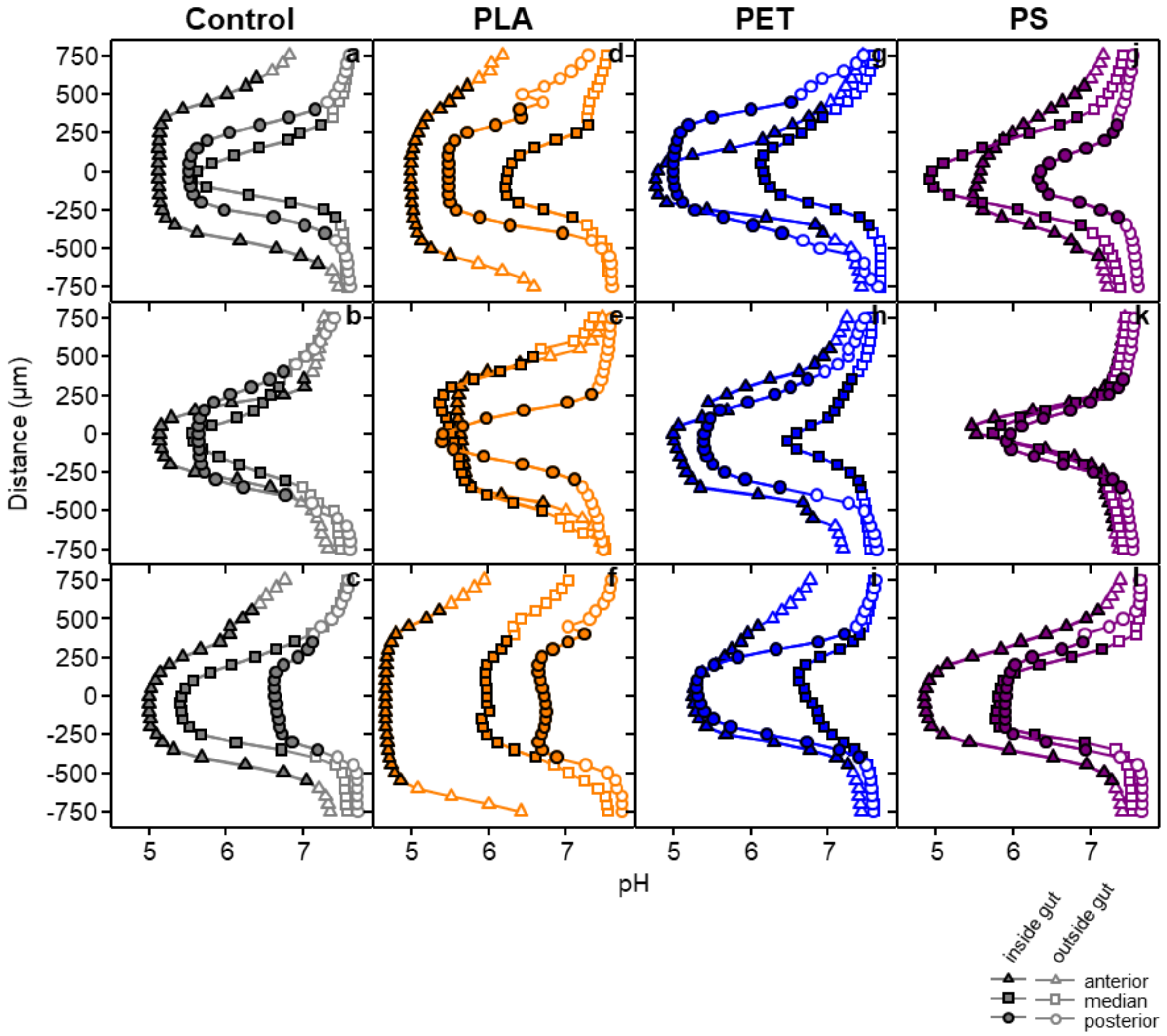


**Fig. S5: Radial pH profiles of woodlice guts.** The woodlice were fed with food containing no microplastic particles (control; **a**, **b**, **c**) or 5% PLA (**d**, **e**, **f**), PET (**g**, **h**, **i**) or PS (**j**, **k**, **l**) for six days prior gut extraction and subsequent embedding in agarose and microsensor measurements. For each gut, profiles were recorded from the anterior, median and posterior. Closed and open symbols represent measured concentrations inside and outside (in agarose) the guts, respectively. Each panel with the same color represents one of three replicate guts per treatment. The distance of 0 µm indicates the center of the tube-like gut.

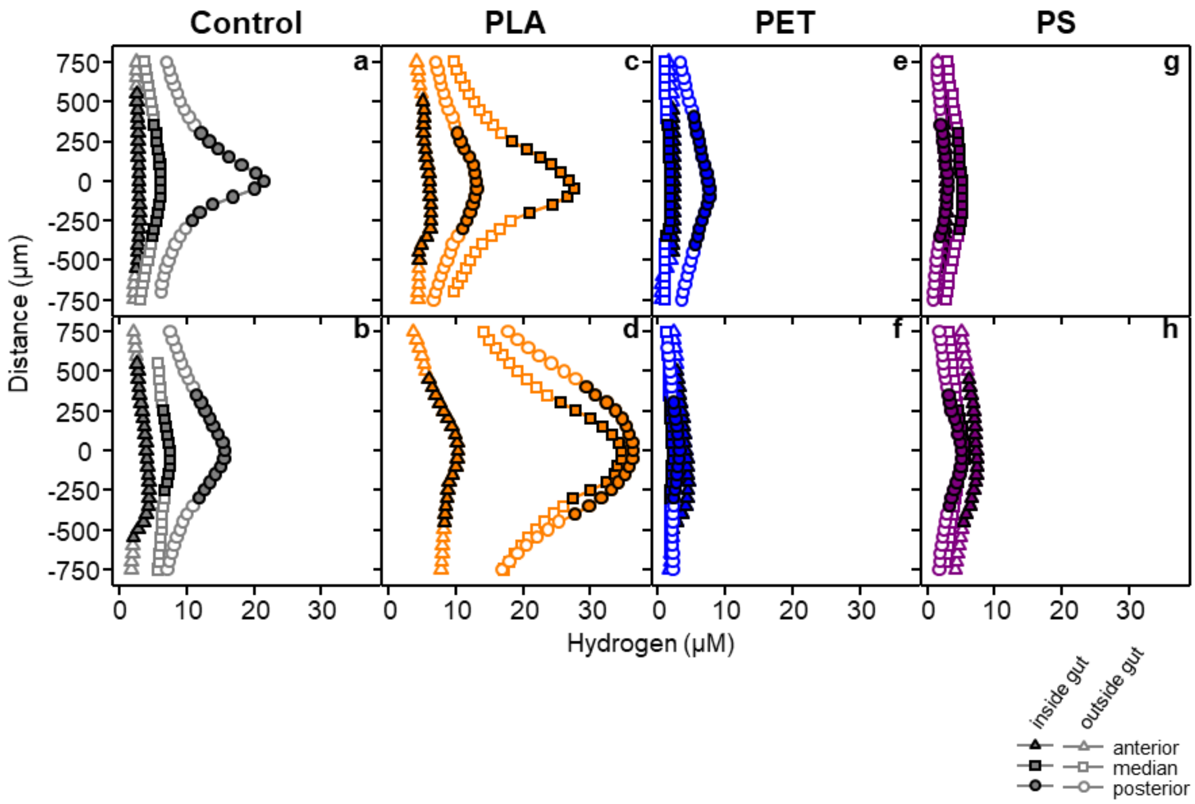


**Fig. S6: Radial hydrogen profiles of woodlice guts in addition to the representative profiles displayed in the main text.** The woodlice were fed with food containing no microplastic particles (control; **a**, **b**) or 5% PLA (**c**, **d**), PET (**e**, **f**) or PS (**g**, **h**) for six days prior gut extraction and subsequent embedding in agarose and microsensor measurements. For each gut, profiles were recorded from the anterior, median and posterior. Closed and open symbols represent measured concentrations inside and outside (in agarose) the guts, respectively. Each panel with the same color represents one of three replicate guts per treatment. The distance of 0 µm indicates the center of the tube-like gut.

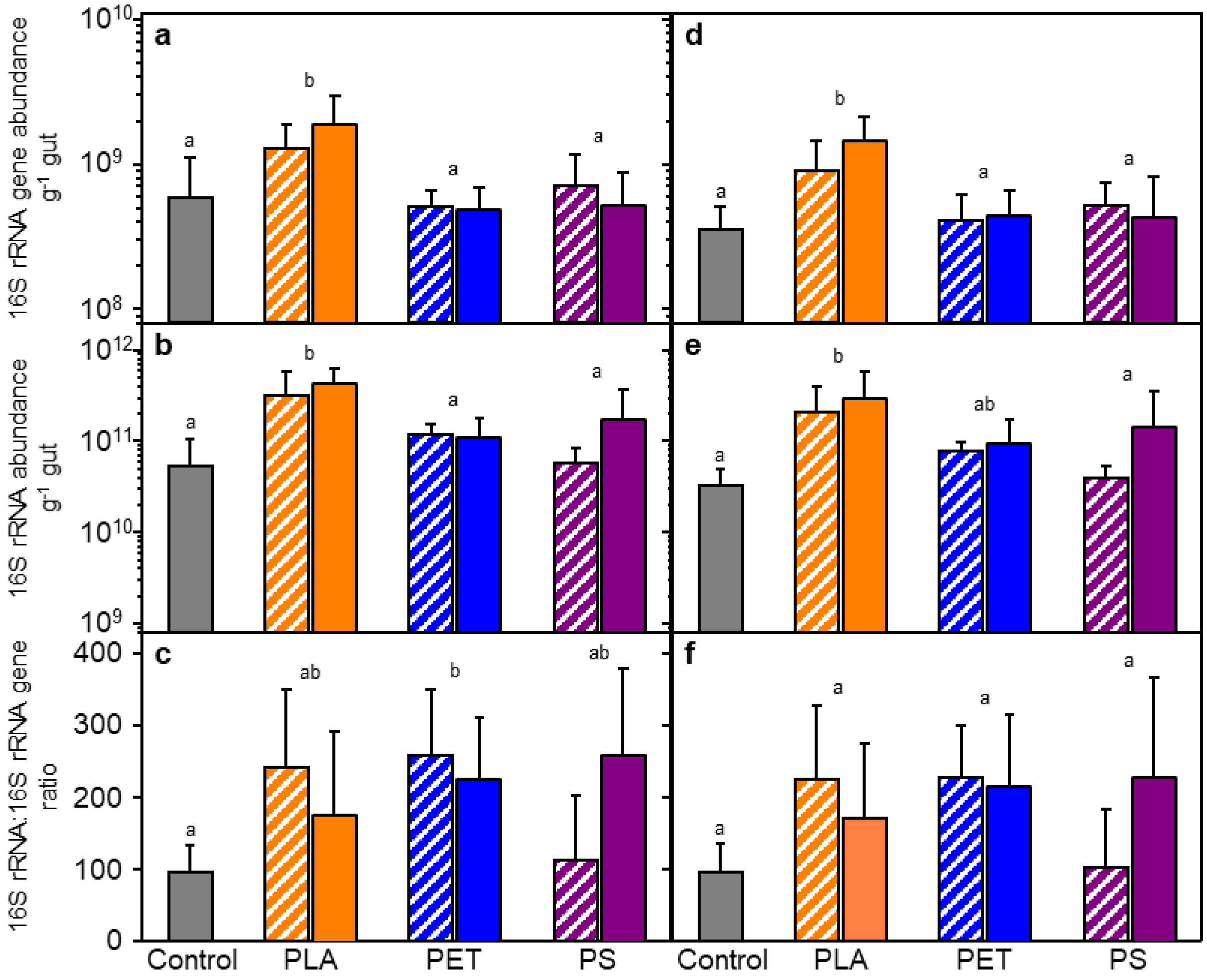


**Fig. S7: Comparison of uncorrected and corrected bacterial 16S rRNA gene (a, d) and 16S rRNA (b, e) abundances and ratios (c, f) in the gut of *P. scaber* exposed to different MP-food.** Nucleic acid extracts were derived from the guts of isopods exposed to control- or 2.5%-MP- (stripped) or 5%-MP- (no pattern) food pellets. Sequencing analysis revealed noticeable occurrences of *P. scaber* specific endosymbionts (see supplementary Fig. S5). The abundance data was corrected for the proportions of these endosymbionts (**d, e, f**). Uncorrected data is~~n~~ displayed for comparison (**a, b, c**). Occurrences of such endosymbionts were marginal in the food pellets and therefore correction was not required. Means and standard deviations of five replicates are plotted. Statistical analysis revealed no effect of the concentration of MP applied and therefore, significant differences in means indicated by different lower letters above the bars are related to the MP treatment regardless the dosage.

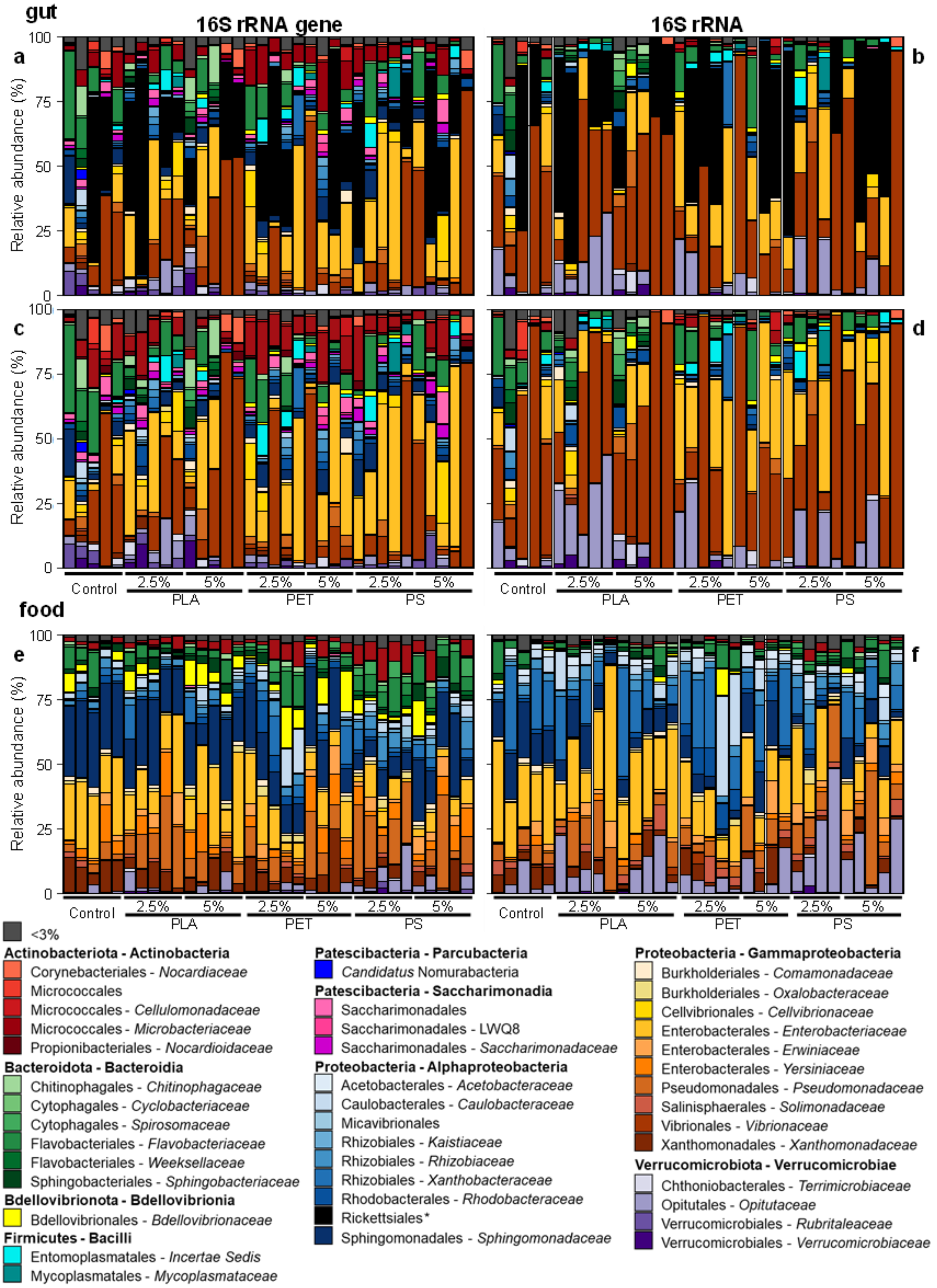


**Fig. S8: Composition of the bacterial community on 16S rRNA gene and 16S RNA level.** Relative abundances of taxa (genera, if applicable; each with at least 3% abundance in one sample) based on 16S rRNA genes (**a, c, e**) and 16S rRNA (**b, d, f**) in the gut of isopods fed with control- or MP-containing food pellets (**a-d**), and in the food pellets (**e, f**). Relative abundances of the communities are shown with (**a, b**) and without (**c, d**) host specific endosymbionts (Rickettsiales - *Wolbachia* and *Candidatus* Hepatincola – in black and indicated by * in the legend). Occurrences of such endosymbionts were marginal in the food pellets.

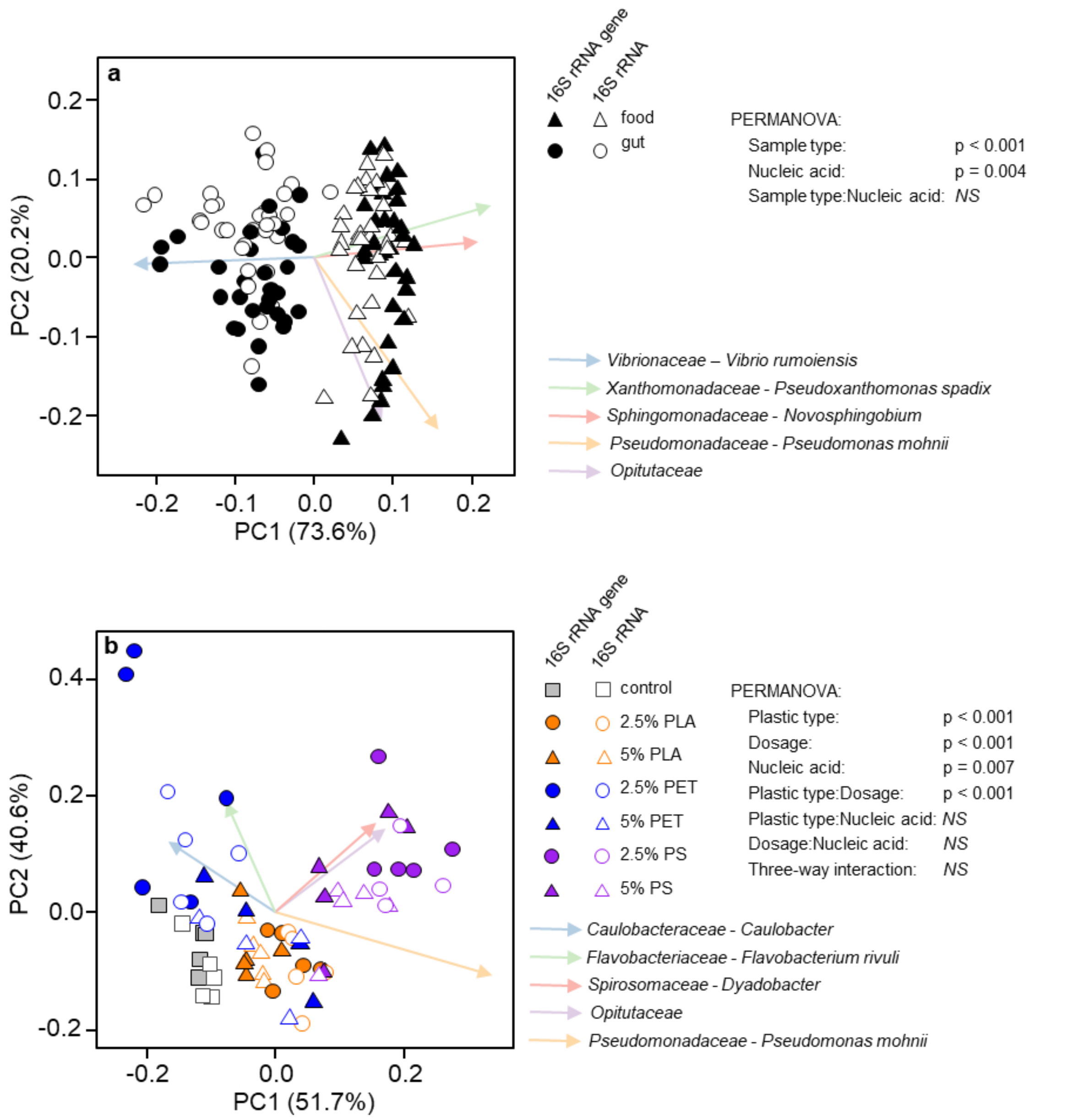


**Fig. S9: Beta diversity of bacterial communities on 16S rRNA gene and 16S RNA level.** PCoA plots are based on Aitchison distance matrixes derived from analyses of the 16S rRNA genes and 16S rRNA of the whole data set (**a**) and of the food communities only (**b**). Results of the PERMANOVA analyses are given. Arrows represent ASVs assigned on family and genus/species level (if applicable) that were highly correlated with the separation of samples.

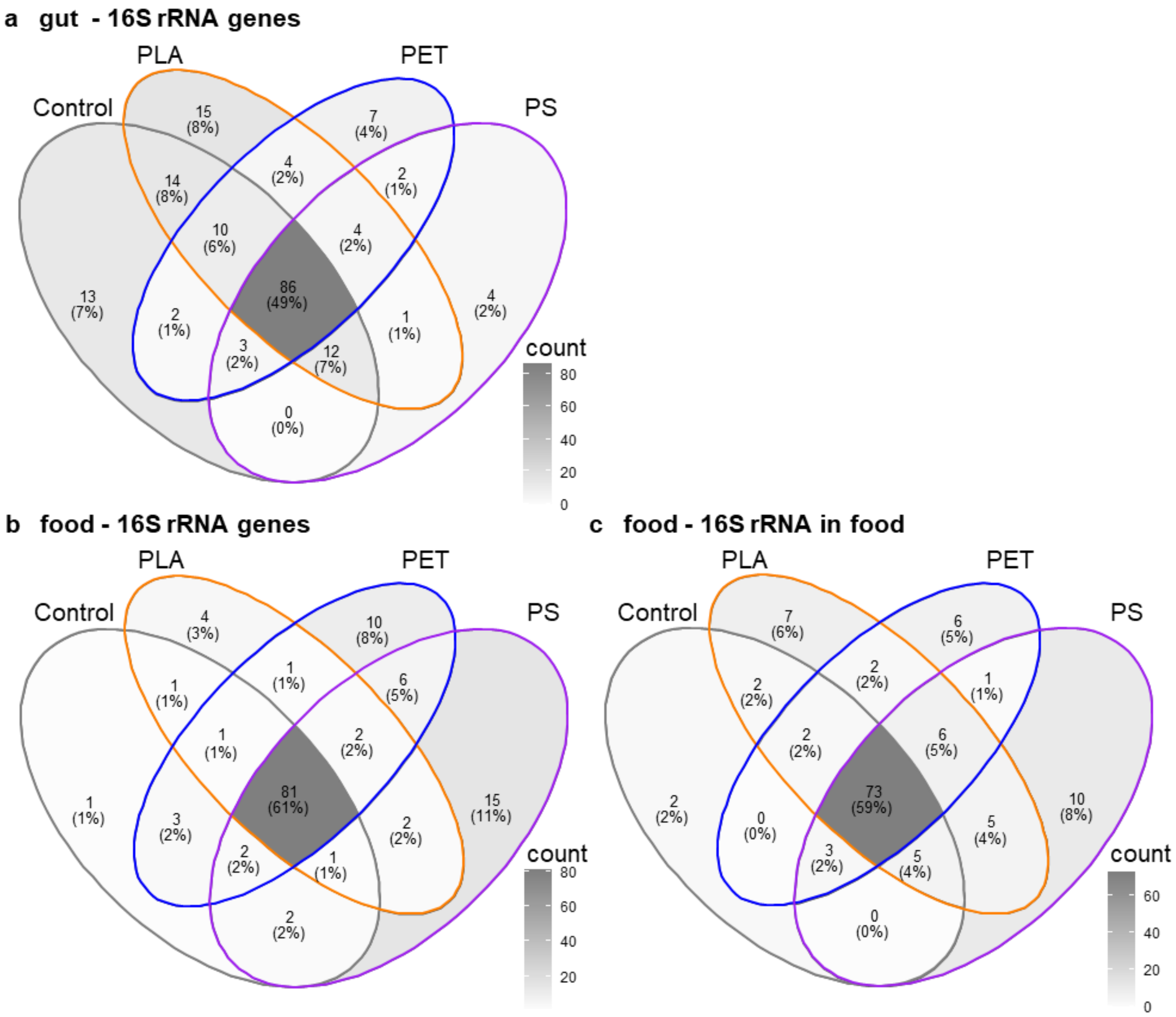


**Fig. S10: Shared and unique numbers and proportions of taxa among isopod guts (a) and food (b, c) pellets on 16S rRNA gene (a, b) and 16S rRNA (c) level.** The Venn diagram for the gut communities on 16S rRNA level is presented in the main text (Fig. 5b). Only taxa on genus level (if applicable) that occurred in at least 30% of the replicates were included for the calculation of the Venn diagrams.

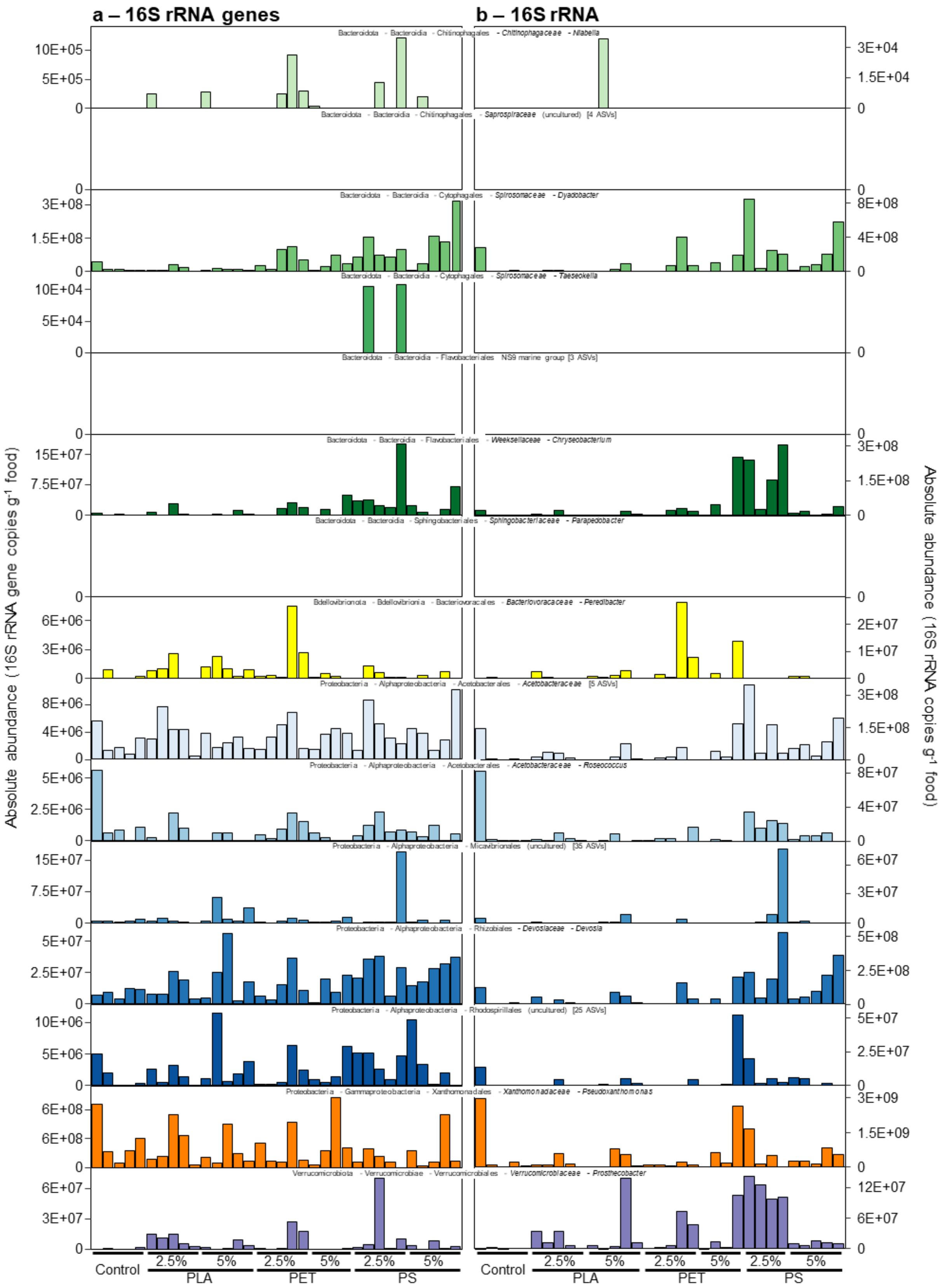


**Fig. S11: Absolute abundance of taxa in the food that were identified as active indicators in the guts of isopods fed with PLA-food.** The abundance of these taxa in isopod guts are shown in Fig. 6a with similar color coding. The relative abundances of taxa were normalized with the total 16S rRNA gene and 16S rRNA abundance derived from qPCR analysis.

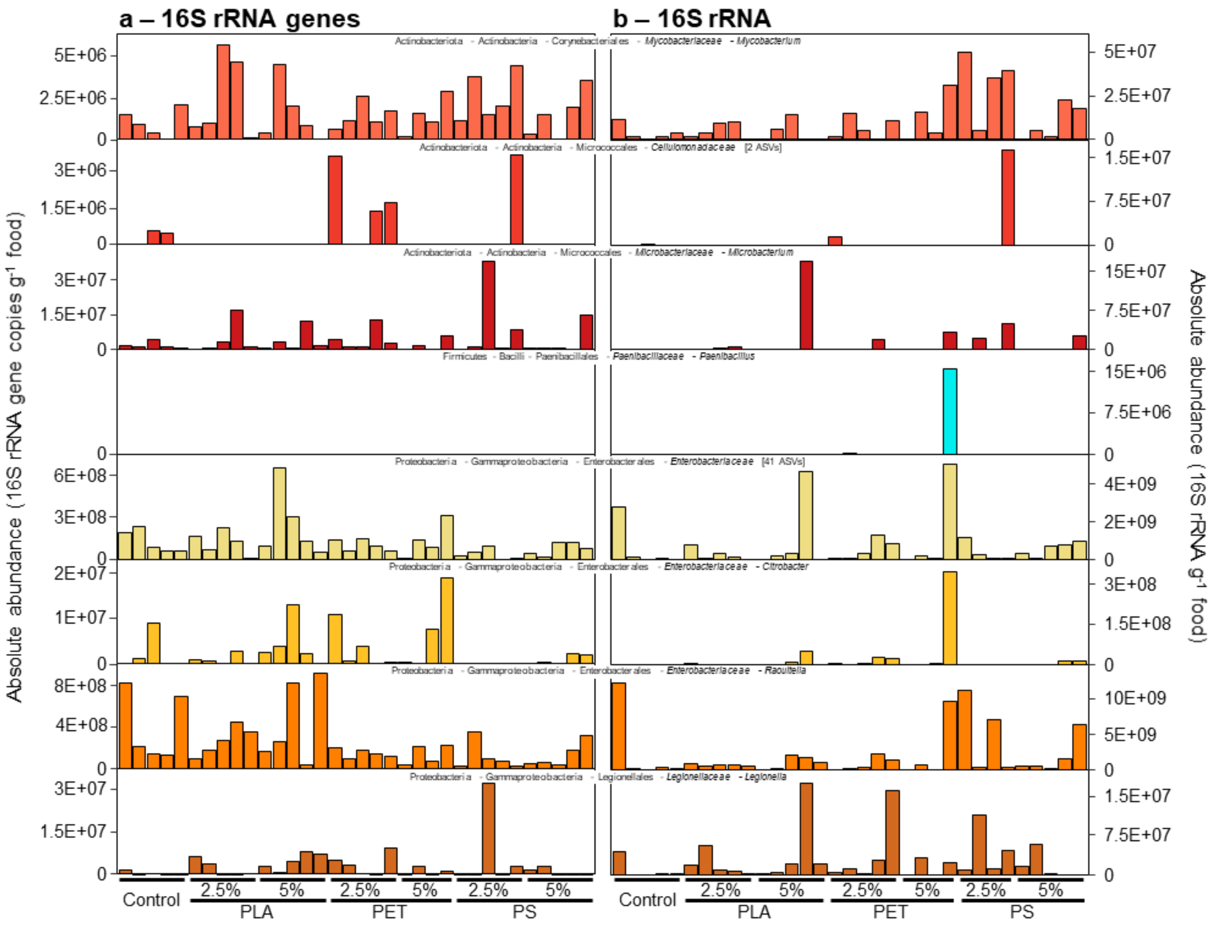


**Fig. S12: Absolute abundance of taxa in the food that were identified as active indicators in the guts of isopods fed with PET-food.** The abundance of these taxa in isopod guts are shown in Fig. 6b with similar color coding. The relative abundances of taxa were normalized with the total 16S rRNA gene and 16S rRNA abundance derived from qPCR analysis.

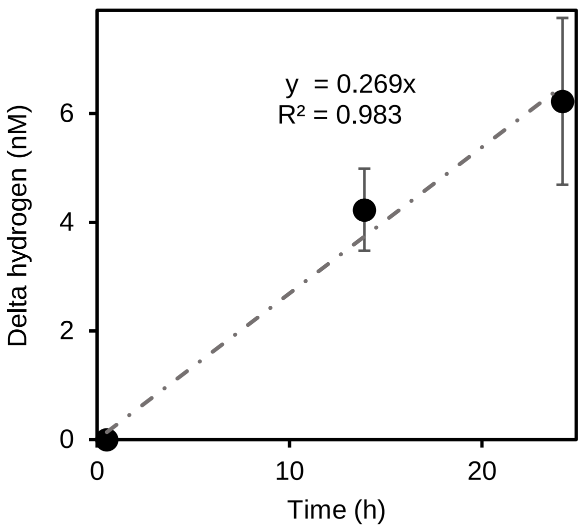


**Fig. S13: Lactate-dependent production of molecular hydrogen by cell suspensions of *Escherichia coli* (*Enterobacteriaceae*) with lactate as carbon source and electron donor.** Hydrogen production of controls incubated under the same conditions in medium without lactate was subtracted. Means and standard deviations of triplicate incubations are plotted. The equation of the regression line and the coefficient of determination ($R^2$) are provided.

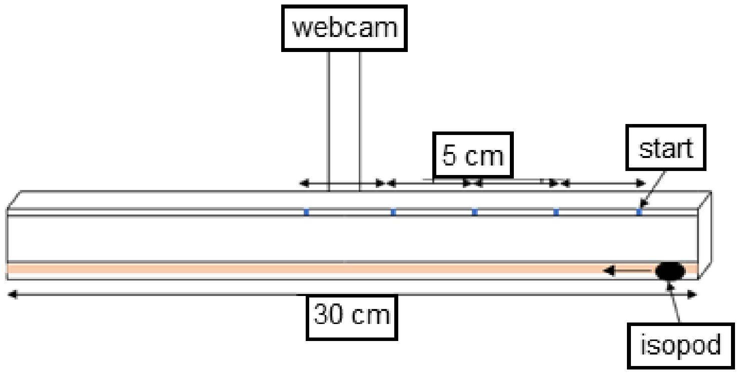


**Fig. S14: Schematic layout of the racetrack used for the evaluation of the locomotor activity of *P. scaber.*** The isopods were placed on a 30 cm racetrack with four marks every 5 cm. Records were taken with a webcam attached over the racetrack.

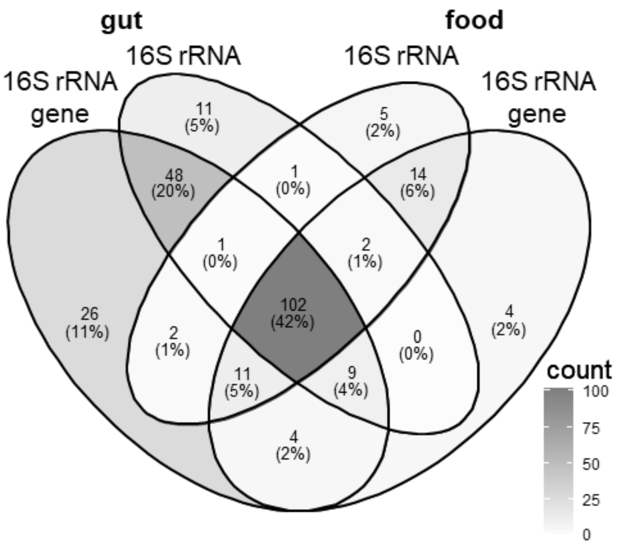


**Fig. S15: Shared and unique numbers and proportions of taxa among guts and food pellets on 16S rRNA gene and 16S rRNA level**. Only taxa on genus level (if applicable) that occur in at 30% of the replicates were included for the calculation of the Venn diagrams. The scale indicates the count numbers of taxa in correlation with the intensity of the shading.
